# Converting Lysosomes into Photothermal Organelles Enables Nanoparticle-Free Tumor Ablation via Intracellular Vapor Bubbles

**DOI:** 10.64898/2026.01.23.698471

**Authors:** Tao Lu, Cristina Muntean, Félix Sauvage, Deep Punj, Herlinde De Keersmaecker, Femke Baeke, Riet De Rycke, Kelly Lemeire, Philippe Tummers, Weiran Li, Chloë De Clercq, Bernd Vanmeerhaeghe, Kaat Durinck, Olivier De Wever, Katrien Remaut, Kevin Braeckmans, Stefaan C. De Smedt, Koen Raemdonck

## Abstract

Photothermal nanomaterials enable precise tumor ablation but face limitations in biodistribution, tissue penetration, toxicity, and biodegradability. Here, we present a unique concept for nanoparticle-free photothermal therapy based on the lysosomal entrapment of cationic amphiphilic small molecular dyes for spatially controlled vapor bubble (VB)-mediated tumor cell ablation. This strategy, which exploits a universal biological and physical effect, employs intracellular pH gradients for extensive local dye enrichment in acidified organelles, transforming them into transient endogenous nanosized photothermal reactors for subsequent light activation. Using sunitinib, a clinically approved lysosomotropic anticancer drug, and the commercially available dye LysoTracker™ Deep Red, lacking intrinsic anticancer activity, we demonstrate pulsed laser-induced VB formation specifically from dye-loaded lysosomes, leading to selective photomechanical disruption of various cancer cell models across 2D cultures, 3D spheroids, patient-derived neuroblastoma tumoroids and tumor fragments from an ovarian carcinoma patient. This approach allows precise, low-fluence and wavelength-tunable cancer tissue ablation without the need for synthetic photoresponsive nanoparticles.

---

Nanoparticle-mediated photothermal therapy (PTT) has emerged as a promising strategy for killing accessible cancer cells using light.^1, 2, 3^ Compared to traditional anticancer treatments, phototherapies typically offer the advantage of high spatiotemporal precision.^4, 5, 6, 7^ However, conventional PTT induces tumor cell death by heat induction, which may result in excessive heat dissipation and potential damage to healthy surrounding tissue.^8^ Laser-induced vapor bubble (VB) formation, mediated by pulsed laser irradiation of photothermal nanoparticles such as gold nanoparticles (AuNPs), iron oxide nanoparticles (IONPs), and polydopamine nanoparticles (PDNPs), has therefore attracted increasing attention.^9^ Upon exposure to femto-, pico- or nanosecond laser pulses at specific wavelengths, such nanoparticles rapidly heat up, causing the surrounding water to vaporize and form VBs, thus mitigating excess heat accumulation and dissipation.^10, 11, 12^ VBs can persist for tens to hundreds of nanoseconds before collapsing, thereby generating mechanical forces capable of locally disrupting targeted cells or tissues.^13^

However, despite these advances, inefficient and heterogeneous intratumoral penetration of nanoparticles remains a central obstacle to effective tumor therapy.^14, 15, 16, 17^ Following systemic administration, nanoparticle transport is strongly constrained by abnormal tumor vasculature, elevated interstitial fluid pressure, and dense tumor stroma, resulting in preferential perivascular accumulation and poor access to deeper tumor regions.^18, 19, 20^ These intrinsic transport barriers are highly variable across tumor types and patients, making it difficult to predict or control intratumoral nanoparticle penetration. This limits the translational impact of many nanoparticle-based strategies across both therapeutic and imaging modalities.^21, 22^ Even localized intratumoral injection into accessible tumors often results in shallow penetration depths and non-uniform tissue coverage, while off-target accumulation and poor biodegradability continue to challenge clinical translation and long-term safety.^23, 24, 25, 26^

To overcome these limitations, small molecules generally offer superior tissue distribution compared to nanoparticles. However, freely dissolved small molecular dyes typically fail to act as efficient photothermal reactors for VB induction due to suboptimal photophysical properties.^27, 28^ Cationic amphiphilic drugs (CADs) are a heterogenous class of weakly basic molecules with high apparent volumes of distribution *in vivo*.^29^ Moreover, due to the pH partition effect and tumor acidosis, CADs are even more likely to accumulate in solid tumors.^30^ Intracellularly, the pH gradient between the near-neutral cytosol (pH ∼7.2) and the acidic lumen of lysosomes (pH ∼4.5) may cause an up to 1000-fold drug accumulation in the lysosomal compartment dependent on the CAD’s pKa.^31, 32^ In this work, we propose to exploit this lysosomotropic behaviour as a nanoparticle-free strategy for photothermal tumor cell ablation. More specifically, upon treating cancer cells with light-sensitive CADs, we hypothesize the spontaneous and transient conversion of drug-loaded lysosomes into photothermal organelles for intracellular VB generation and tumor cell eradication. The latter builds on our recent work demonstrating melanoma cell ablation via VBs generated from intracellular melanosomes under nanosecond laser irradiation.^33^ The proposed strategy extends this concept to non-pigmented cancer cells, aiming to make the approach broadly applicable.

As a proof-of-concept, we first repurposed the clinically approved cationic amphiphilic tyrosine kinase inhibitor sunitinib to induce intracellular laser-triggered VBs from lysosomal organelles. To demonstrate the broader applicability of the concept beyond repurposed drugs, lysosomal VB formation was also evaluated using LysoTracker™ Deep Red (LDR), a well-known cationic amphiphilic dye that accumulates in lysosomes via a similar mechanism. Exposure of sunitinib- or LDR-treated cancer cells to nanosecond laser pulses enabled effective and spatial-selective VB-mediated cell killing and tumor ablation in various 2D and 3D tumor cell models, including patient-derived neuroblastoma tumoroids and tumor fragments from a low-grade serous ovarian cancer (LGSOC) patient.

Taken together, our findings highlight the potential of lysosomotropic cationic amphiphilic photosensitizers as a general platform for lysosome-mediated intracellular VB formation, enabling nanoparticle-free tumor cell eradication with high spatial precision.

## Accumulation and intracellular localization of sunitinib

Owing to its physicochemical properties as a hydrophobic weak base (logP: 5.2; pKa: 9.0, DrugBank database), sunitinib is expected to spontaneously accumulate in the lysosomes of cancer cells (Fig. 1a). Sunitinib exhibits strong absorbance in the 340-490 nm range and fluoresces around 540 nm, making it inherently suitable for light activation (Supplementary Fig. 1).^34, 35^ Detection of sunitinib’s fluorescence by flow cytometry indicated that intracellular sunitinib levels in three different cancer models (i.e., HeLa cells, Jurkat cells and HT1080 cells) rise in a concentration-dependent manner, with all cells having acquired the compound at concentrations ≥ 2.5 µM and 6 h of incubation (Supplementary Fig. 2a). The relative mean fluorescence intensity (rMFI) also revealed a time-dependent intracellular accumulation of sunitinib, with 24 h incubation showing highest sunitinib fluorescence in the cells (Fig. 1b and Supplementary Fig. 3). Confocal microscopy imaging of the intracellular sunitinib distribution showed a clear co-localization of the drug with LDR-stained organelles (Fig. 1d and Supplementary Fig. 4), corroborating its lysosomotropic behaviour.^36^ Moreover, sunitinib exposure increased the intracellular lysosomal density, indicated by both a heightened rMFI (Fig. 1c) and LDR fluorescence area per cell (Fig. 1e and Supplementary Fig. 5). The latter may result from both sunitinib-induced lysosomal swelling as well as *de novo* lysosomal biogenesis, as generally described for lysosomotropic CADs.^37^ Additionally, sunitinib exposure increased the side scatter (SSC-A) signal, a parameter indicative of lysosomal perturbation by CADs, in a time- and concentration-dependent manner (Supplementary Fig. 2b).^38, 39^ Altogether, exposing cancer cells to sunitinib shows the typical hallmarks of lysosomotropism.

**Fig. 1.**
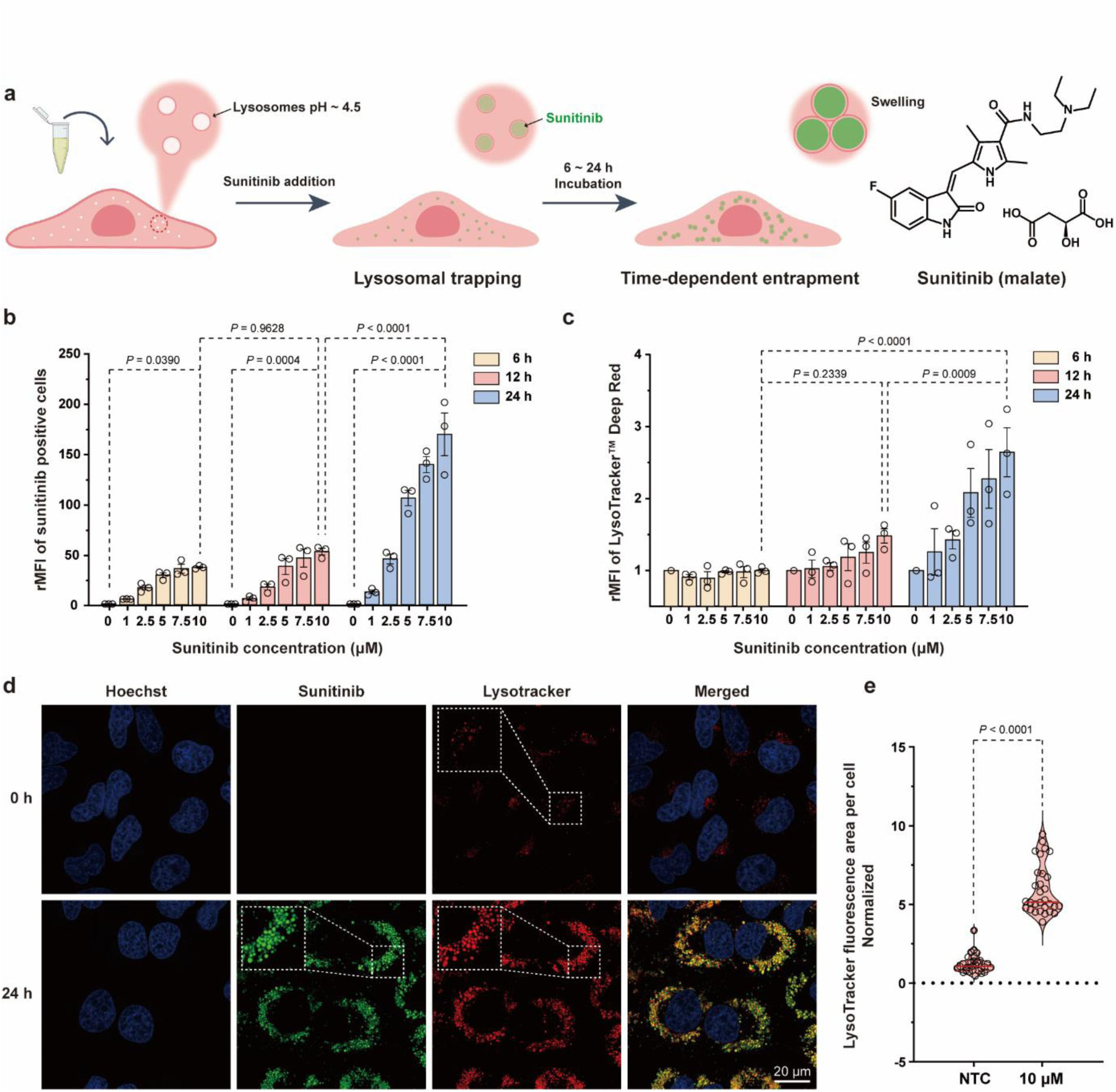
Characterization of sunitinib accumulation in HeLa cells. **(a)** Schematic of intracellular sunitinib accumulation and lysosomal sequestration. **(b)** Relative mean fluorescence intensity (rMFI) of cells accumulating sunitinib. **(c)** Relative increase in Lysotracker™ Deep Red (LDR) staining of HeLa cells as a function of sunitinib concentration and incubation time. Data are presented as mean ± SEM (N=3, n=3). **(d)** Sunitinib accumulation (5 µM, 24 h) and co-localization with LDR staining. **(e)** Quantification of LDR fluorescence area per cell (normalized). Data are presented as median ± interquartile range (N=3, n=10).

## Pulsed laser irradiation of lysosomal accumulated sunitinib triggers VB formation

Considering the optical properties of sunitinib and its high degree of lysosomal accumulation, we hypothesized that sunitinib-loaded lysosomes could serve as endogenous intracellular photothermal organelles. Upon nanosecond pulsed laser irradiation, the photothermal conversion of the lysosomes may lead to rapid heating, potentially inducing the formation of intracellular VBs (Fig. 2a). To verify this, HeLa cells, A549 cells, HT1080 cells and Jurkat cells were treated with 10 µM sunitinib during 24 h and successively irradiated with a nanosecond pulsed laser (480 nm) with increasing laser fluence (*i.e.*, energy applied per unit area, J cm^-2^). Dark-field microscopy clearly revealed the emergence of intracellular VBs via detection of the extensive light scattering produced during their lifetime (Fig. 2b,d and Supplementary Movie 1). In contrast, laser irradiation of sunitinib in solution (up to 2 mM) did not evoke VBs under identical conditions (Supplementary Movie 2), indicating that accumulation of sunitinib in the lysosomes is required. Quantification of the number of VBs formed within a defined laser-irradiated area as a function of the laser pulse fluence allowed to calculate the VB generation threshold (Fig. 2c,e), which is defined as the fluence of a single laser pulse at which VBs are formed with 90% probability. Based on the Boltzmann fitting parameters, the VB formation process can be categorized into three distinct regimes. The “Sparse” regime, spanning from the baseline to the inflection point, is characterized by a low and stochastic VB count. This is followed by the “Moderate” regime, defined between the inflection point and the 90% threshold, where the VB population increases significantly and non-linearly per pulse before reaching the saturation plateau. The VB threshold exhibited minor variation among different adherent cell types. For instance, HeLa cells and A549 cells displayed a threshold of 1.0 J cm^-2^, whereas Jurkat cells formed VBs at a substantially lower laser fluence of 0.8 J cm^-2^ (Supplementary Movie 3). A plausible explanation for the latter is that Jurkat cells tend to form clusters, allowing multiple sunitinib-loaded cells to contribute cooperatively to localized heat generation, facilitating VB formation at lower energy inputs. In support of this, resuspending Jurkat cells as a loose monolayer increased the VB threshold to 3.0 J cm^-2^ (Supplementary Fig. 6). Jurkat monolayers requiring substantially higher laser fluences than 2D cultures of either HeLa or A549 cells can be explained by the markedly lower sunitinib accumulation (Supplementary Fig. 2c and Supplementary Fig. 3b). The latter likely limits localized intra-lysosomal heating upon laser irradiation, thereby raising the energy threshold for VB formation. Furthermore, according to the pH partition theory, a higher initial extracellular sunitinib dose will lead to higher intra-lysosomal drug concentrations, thus increasing the photo-responsiveness of the organelles and lowering the VB threshold (Supplementary Fig. 7a,b). Hence, in cells treated with 20 µM sunitinib, VB formation was already detectable at a laser fluence as low as 0.3 J cm^-2^ (Supplementary Fig. 7d). HeLa cells incubated with 10 µM sunitinib undergo immediate and complete ablation upon a single laser pulse, while at a lower concentration of 5 µM no visible change in cellular morphology was observed (Supplementary Movie 4). The applied laser fluence of 1.7 J cm^-2^ is above the threshold for 10 µM sunitinib, while it fails to generate VBs in cells exposed to 5 µM sunitinib, supporting that cell disruption is caused by VBs. At the highest sunitinib concentration of 20 µM, even the formation of large microbubbles was observed, presumably formed by coalescence of multiple VBs in the irradiated area (Fig. 2f and Supplementary Fig. 7d). After laser application, the cells in this illuminated spot were again clearly ablated (Fig. 2g and Supplementary Movie 5). Comparable results were also obtained for the A549 cell type (Supplementary Fig. 8). In summary, incubation of a variety of cancer cell lines with sunitinib can induce intracellular VBs from drug-loaded lysosomal compartments upon pulsed laser irradiation, dependent on the applied sunitinib concentration, incubation time and laser fluence.

**Fig. 2.**
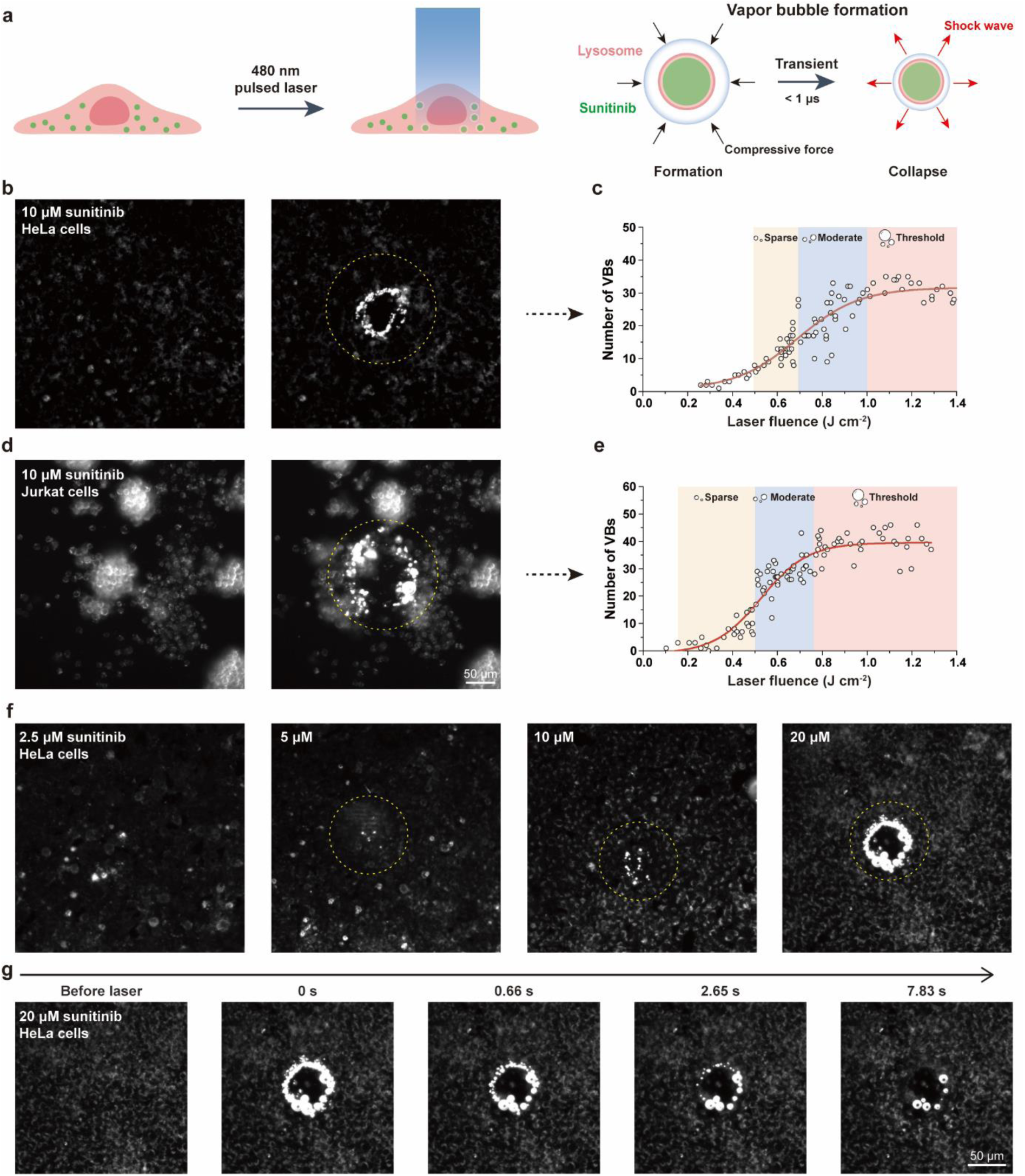
Detection of vapor bubbles (VBs) in sunitinib-exposed cells upon nanosecond pulsed laser irradiation. **(a)** Schematic of laser irradiation of cancer cells with sunitinib-loaded lysosomes to produce intracellular VBs, illustrating the VB formation and collapse from sunitinib-loaded lysosomes. **(b)** Dark-field microscopy images of HeLa cells before and during laser irradiation for visual detection of intracellular VBs, and **(c)** the corresponding quantification of the intracellular VB formation threshold. **(d)** Dark-field microscopy images of Jurkat cells before and during laser irradiation and **(e)** the corresponding intracellular VB formation threshold. The thresholds were determined for HeLa and Jurkat cells treated with 10 µM sunitinib for 24 h. **(f)** Dark-field microscopy images of HeLa cells incubated with different concentrations of sunitinib (2.5-20 μM) and exposed to pulsed laser irradiation at a fluence of 1.0 J cm^-2^. **(g)** Time-resolved dark-field microscopy images capturing the formation and collapse of VBs in HeLa cells incubated with 20 μM sunitinib following a single nanosecond laser pulse. The sequence illustrates the full dynamic process of intracellular VB generation and subsequent collapse.

## Cell killing induced by intracellular VB formation upon pulsed laser irradiation of sunitinib-treated cancer cells

Significant laser-induced cell death was observed across all sunitinib concentrations tested, in contrast to laser treatment alone (Fig. 3a). Near complete HeLa cell eradication was achieved at 10 µM sunitinib and 1.8 J cm^-2^ laser pulses, conditions leading to extensive VB formation and photomechanical cell ablation within the irradiated region. Higher laser fluences and sunitinib concentrations likely induce more damage to the cell membrane that cannot be easily repaired, thereby causing greater cytotoxicity (Fig. 3b).^40^

**Fig. 3.**
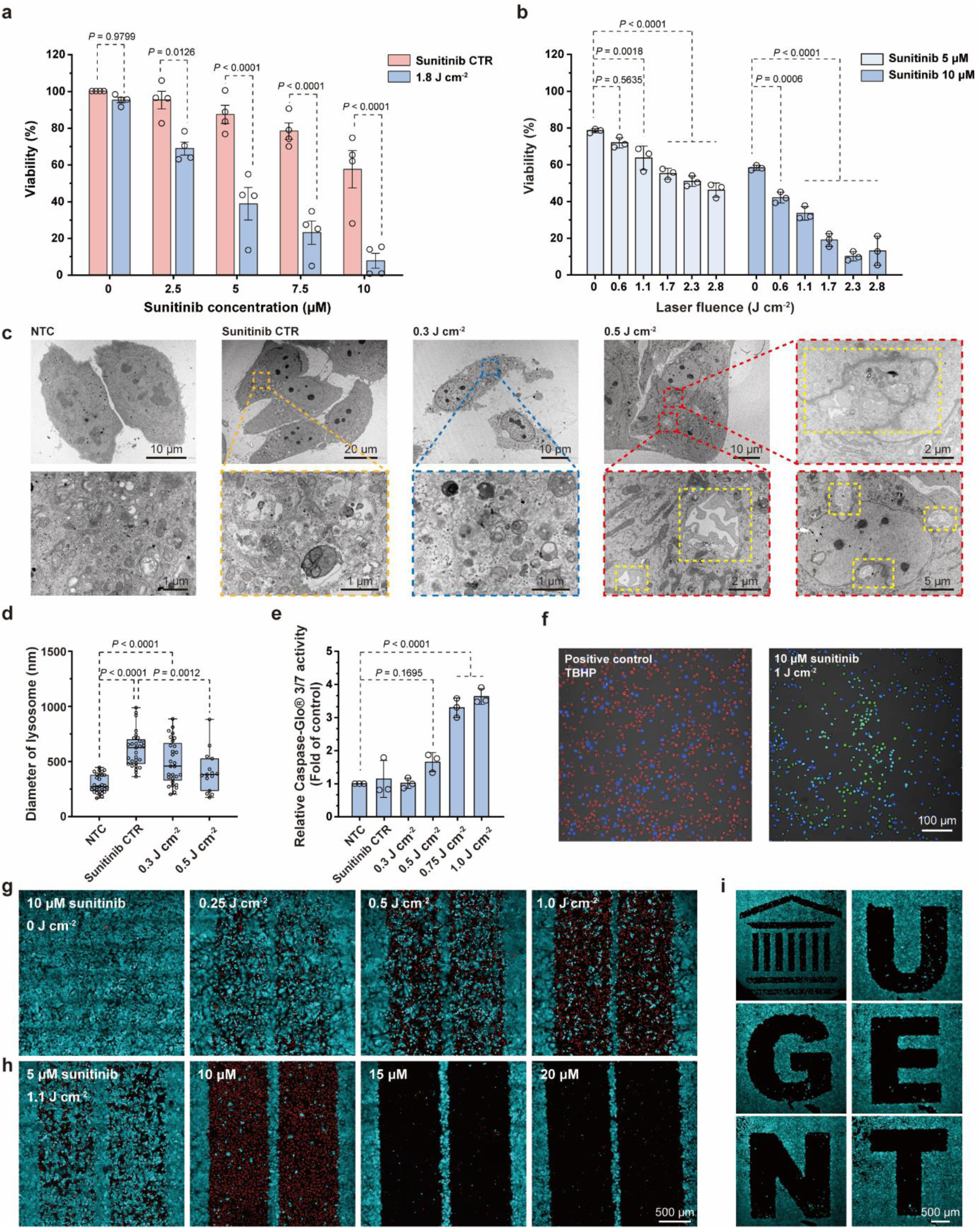
Pulsed laser-induced cell killing of sunitinib-exposed cells. **(a)** Decrease in cell viability after laser treatment (1.8 J cm^-2^) applied on HeLa cells incubated with sunitinib for 24 h at concentrations of 0-10 µM. **(b)** Cell viability of HeLa cells treated with sunitinib (5 and 10 µM) decreased with increasing laser fluence. **(c)** Transmission electron microscopy (TEM) images of HeLa cells incubated with 10 µM sunitinib for 24 h for sunitinib only control (no laser, sunitinib CTR), low-fluence irradiation (0.3 J cm^-2^), and VB-generating fluence (0.5 J cm^-2^). Low-magnification images (top) depict overall cellular morphology, while corresponding high-magnification views (bottom) focus on lysosomal structures. Yellow dashed boxes mark cytoplasmic voids observed after VB formation. **(d)** Quantification of average lysosome diameter in HeLa cells based on TEM analysis after different treatments. **(e)** Quantification of caspase-3/7 activation in HeLa cells incubated with 10 µM sunitinib for 24 h, followed by laser irradiation at increasing fluences (0.3-1.0 J cm^-2^). Caspase activity was measured using the Caspase-Glo® 3/7 assay and normalized to the non-treated control (NTC) group. **(f)** Assessment of intracellular reactive oxygen species (ROS) levels in Jurkat cells using the CellROX™ Deep Red assay following nanosecond pulsed laser irradiation. **(g)** Confocal microscopy images of HeLa cells treated with 10 µM sunitinib after scanning with different laser fluences in a pre-defined line pattern and **(h)** images of cells treated with different concentrations of sunitinib (5-20 µM) using a fixed laser fluence (1.1 J cm^-2^). **(i)** Confocal microscopy images of cells scanning in a pre-defined text pattern (10 µM, 1.1 J cm^-2^). Data are presented as mean ± SEM (N=3, n≥3).

To investigate the subcellular events associated with VB formation, we next performed transmission electron microscopy (TEM) on HeLa cells following sunitinib incubation and laser irradiation (Fig. 3c). In line with earlier observations, 10 µM sunitinib-treated cells exhibited pronounced lysosomal swelling. When cells were irradiated with laser fluences below the VB threshold (e.g., 0.3 J cm^-2^), serving as a heating-only control, no obvious morphological changes were observed. At a laser fluence of 0.5 J cm^-2^, corresponding to the onset of VB formation at the given sunitinib concentration, distinct cytoplasmic voids emerged, suggestive of sites of VB formation and subsequent collapse. The latter coincides with a significant decrease in the number of intact lysosomes, and a reduction in their size, indicative of selective rupture of the larger lysosomes or VB-induced lysosomal fragmentation (Fig. 3d). In some images, these voids appeared to impinge upon or even disrupt the nuclear envelope. Also, cells irradiated at 0.5 J cm^-2^ frequently exhibited elongated mitochondria, a well-established morphological hallmark of cellular stress responses.^41, 42, 43, 44, 45^ Additionally, a Caspase-Glo® 3/7 assay revealed a clear activation of caspase 3/7, particularly when exceeding the VB threshold (Fig. 3e), indicating that apoptosis is triggered after VB formation. As intracellular reactive oxygen species (ROS) generation is a key cytotoxic mechanism in conventional PDT,^46^ the CellROX™ Deep Red Assay was employed to detect ROS production following sunitinib exposure and laser treatment of Jurkat cells. However, both flow cytometry and confocal fluorescence microscopy revealed negligible ROS generation under the applied conditions (Fig. 3f and Supplementary Fig. 9g).

To explore the spatiotemporal nature of sunitinib-induced photoablation, a laser beam (spot size ∼ 200 µm) was scanned across a monolayer of sunitinib-treated HeLa cells in a pre-defined pattern consisting of two vertical lines, each 1 mm wide and separated by a non-irradiated zone of 100 µm (Supplementary Fig. 9a,b). Necrotic and late apoptotic cells, identified by positive TO-PRO-3 staining, were only found within the irradiated regions, whereas adjacent non-irradiated cells remained viable (*i.e.*, calcein-AM positive), despite sunitinib exposure. These observations demonstrate precise spatial control over laser-induced intracellular VB formation and cell ablation (Fig. 3g). Confocal microscopy further confirmed that the extent of photomechanical cell killing was dependent on both sunitinib concentration and laser fluence. At the highest sunitinib concentration tested, TO-PRO-3 positive cells were no longer detectable within the irradiated zones, indicative of severe cell ablation (Fig. 3h). Comparable results were again obtained using A549 cells (Supplementary Fig. 9d). Cells located in non-irradiated areas retained their proliferation capacity after laser treatment, further verifying the spatial selectivity of the laser treatment (Supplementary Fig. 9e). Furthermore, we demonstrated highly selective cell killing through the creation of customized irradiation patterns, including defined letters and geometric shapes (Fig. 3i).

Finally, the transient effect of drug exposure on subsequent laser-induced cell killing was assessed by replacing sunitinib-containing medium (after 24 h incubation) with fresh culture medium for an additional 72 h prior to laser treatment (Supplementary Fig. 9f). A marked reduction in the laser-induced cell killing effect was observed, likely explained by sunitinib wash-out. Furthermore, laser irradiation at suboptimal wavelength (e.g., 532 nm), which lies outside the absorption band of sunitinib, did neither induce any detectable VB formation nor compromise cell viability (Supplementary Fig. 9h). These findings highlight both the requirement for sufficient intra-lysosomal sunitinib concentration to enable VB-mediated cell ablation as well as the reversible nature of the lysosomal sunitinib sequestration.

## Photoablation of sunitinib-treated tumor spheroids induced by VB formation

Spheroids have been described to more closely mimic the architecture, biological behaviour, and microenvironmental characteristics of human tumors.^47, 48, 49^ HeLa cell spheroids were incubated with different sunitinib concentrations for 24 h, resulting in a dose-dependent intracellular drug accumulation and a marked reduction in spheroid diameter (Fig. 4a and Supplementary Fig. 10a). The latter observation likely correlates with the reduced cell viability within the spheroids, approaching near-complete cell death at a concentration of 30 µM sunitinib (Fig. 4b). Due to the higher cell density and increased light scattering and absorption within 3D spheroids compared to 2D cultures, photoablation was performed using multiple consecutive laser pulses (n = 10) at a fluence of 1.0 J cm^-2^. To visualize the ablation effect more clearly, the laser was targeted at the periphery of the spheroid rather than the core.^50, 51^ Light microscopy enabled real-time imaging of VB formation at the spheroid periphery (Fig. 4d). Furthermore, edge-localized photoablation damage could be visualized under dark-field microscopy (Fig. 4e and Supplementary Movie 6).

**Fig. 4.**
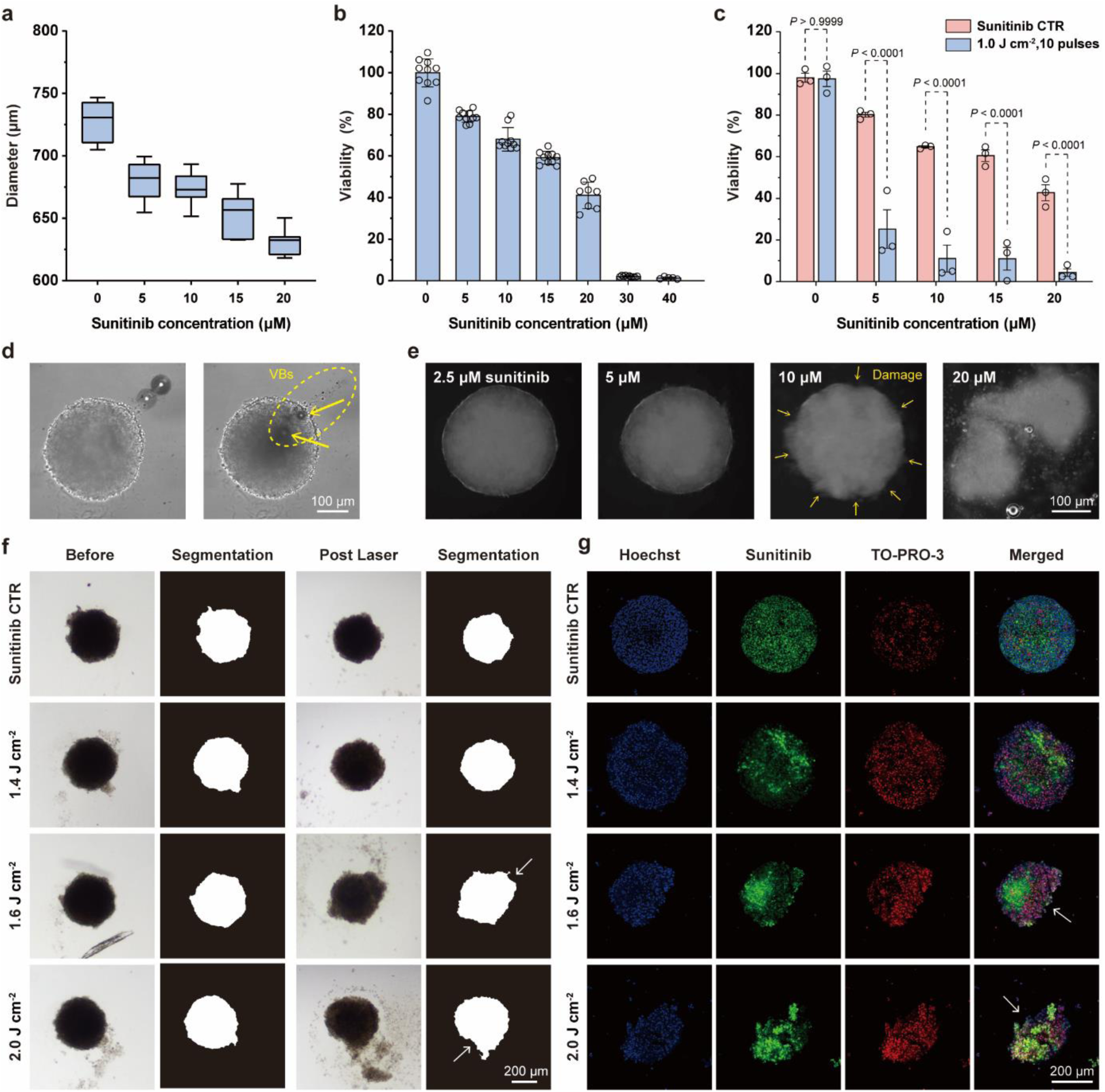
Photoablation of sunitinib-exposed tumor spheroids by pulsed laser irradiation. **(a)** Diameter and **(b)** viability of HeLa spheroids incubated with different concentrations of sunitinib. **(c)** Viability of spheroids before and after laser treatment. Data are presented as mean ± SEM (N≥5, n=3) **(d)** Optical microscopy images showing VBs at the spheroid’s edges as a function of localized laser irradiation. Yellow dashed circles indicate the irradiated region and highlight the structural disruption of the spheroid observed at the moment of VB collapse. Yellow arrows mark prominent VBs formed during laser exposure. **(e)** Dark-field microscopy images showing the photoablation effect of spheroids cultured with mounting concentrations of sunitinib. **(f)** Single time-point analysis of spheroids using bright-field microscopy for morphometric analysis. **(g)** Single time-point analysis of spheroids by confocal fluorescence microscopy. Fluorescent labeling with Hoechst (blue, indicating nuclei) and TO-PRO-3 (red, indicating dead cells) enabled evaluation of fluorescence intensity distribution within the spheroid area. Confocal fluorescence microscopy was performed using Z-stack imaging to further assess the spatial distribution of both markers following laser exposure, shown as maximum intensity projections. Sunitinib CTR, sunitinib only control.

At the lowest sunitinib concentration tested (5 µM), laser treatment resulted in a marked reduction in tumor cell viability from 79% to 25% (Fig. 4c), despite no overt morphological ablation being observed at this dose. By contrast, control spheroids showed no significant decrease in viability upon laser exposure, which confirms that sunitinib accumulation is required. At higher sunitinib concentrations, the effect of laser exposure was further amplified, with <5% viable cells remaining when combining 20 µM sunitinib with laser treatment. This level of cell killing was not reached even with high doses of photothermal NPs (Supplementary Fig. 11c,d), with spheroids demonstrating higher resistance to NPs-induced cell death compared to 2D cultures.

Additional morphometric parameters, such as spheroid circularity and fragmentation, were evaluated by bright-field microscopy combined with automated image segmentation using AnaSP software (Fig. 4f-h).^52^ Upon laser irradiation, especially at fluences of 1.6 J cm^-2^ and above, clear edge defects became visible (white arrows), likely resulting from VB formation and collapse at the spheroid periphery. These structural alterations were further reflected in the quantification of circularity (Supplementary Fig. 10b). Remarkably, although spheroids irradiated at 1.4 J cm^-2^ did not exhibit visible fragmentation, viability analysis indicated over 70% cell death. This suggests that while the majority of cells were non-viable, the spheroid retained its physical integrity, possibly due to the presence of extracellular matrix (ECM) components.^53^ At fluences of 1.6 J cm^-2^ and above, clear spheroid ablation and fragmentation was observed, correlating with increased TO-PRO-3 staining (Fig. 4g). Notably, in spheroids exhibiting fragmentation, the peripheral regions showed markedly elevated TO-PRO-3 fluorescence, indicating that the VB formation caused pronounced cell death at the spheroid edges. This observation highlights the role of photomechanical damage in compromising spheroid integrity and inducing localized ablation, which demonstrates the potent ability to kill sunitinib-loaded tumor cells, even within dense 3D structures.

## Repurposing lysosomotropic dyes for lysosome-assisted intracellular VB formation

Next, we sought to investigate whether other photo-responsive lysosome-targeting dyes might elicit a similar photomechanical response. To validate this, LysoTracker™ Deep Red (LDR), a commercially available probe used for visualizing acidic organelles, was selected (Fig. 5a). Unlike sunitinib, LDR lacks intrinsic anticancer activity and exhibits strong absorption in the 500-600 nm range (Supplementary Fig. 1d). Notably, LDR exhibits a relatively low fluorescence quantum yield (reported to be < 0.4),^54, 55^ indicating that a substantial fraction of the absorbed photon energy is dissipated through non-radiative decay pathways. This enables efficient photothermal conversion required for VB formation. First, HeLa cells were incubated with 0.5 µM or 1 µM LDR for 24 h and subsequently irradiated with a 532 nm nanosecond pulsed laser, using a commercially available benchtop photoporation device (LumiPore™, Trince, Ghent, Belgium).^9, 10, 56^ Clear VB formation was observed, with lower LDR concentrations requiring higher laser fluences (Fig. 5b,c and Supplementary Fig. 12a,b). Similar results were obtained in HT1080 cells (Fig. 5d,e and Supplementary Movie 7). Time-resolved dark-field imaging in both HeLa and HT1080 cells incubated with 1 µM LDR and irradiated at 1.2 J cm^-2^ clearly illustrate the VB lifecycle, demonstrating a VB lifetime within 10 ms (Supplementary Fig. 13a). Cells irradiated at 561 nm, a wavelength closer to the absorption maximum of LDR, generated more VBs compared to 532 nm at a slightly reduced VB threshold (Supplementary Fig. 14). Altogether, these data confirm VB generation in LDR-treated cells following pulsed laser irradiation at a wavelength matching the dye’s absorption spectrum.

**Fig. 5.**
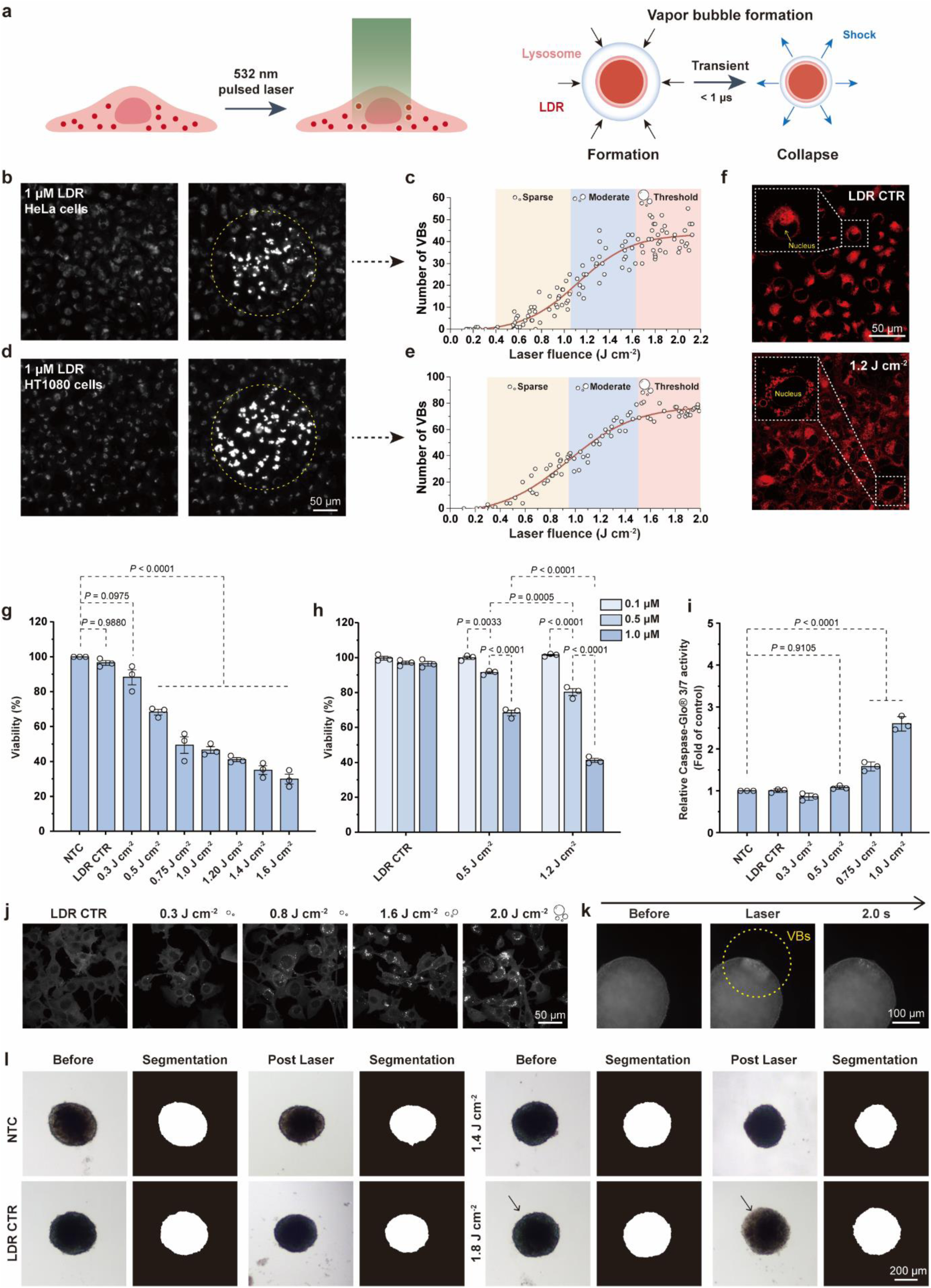
Detection of vapor bubbles (VBs) in LDR-treated cells upon nanosecond pulsed laser irradiation. **(a)** Schematic of pulsed laser irradiation using the LumiPore™ platform of lysosome-stained cells with Lysotracker™ Deep Red (LDR) to produce VBs, illustrating their formation and collapse. **(b-e)** Dark-field microscopy of HeLa and HT1080 cells before and during laser application and visual detection of VBs and VB threshold quantification for cells treated with 1 µM LDR for 24 h. **(f)** Confocal fluorescence microscopy of HeLa cells treated with 1 µM LDR before and after laser irradiation at 1.2 J cm^-2^, illustrating intracellular morphological deformations following VB generation and collapse. **(g)** Cell viability of HT1080 cells treated with LDR (1 μM) with increasing laser fluence. **(h)** Quantification of HT1080 cell viability after incubation with different concentrations of LDR (0.1-1 µM), followed by pulsed laser irradiation at 0.5 J cm^-2^ and 1.2 J cm^-2^. **(i)** Quantification of caspase-3/7 activation (Caspase-Glo® 3/7 assay) in HT1080 cells incubated with 1 µM LDR for 24 h, followed by laser irradiation at increasing fluences (0.3-1.0 J cm^-2^). Caspase activity was normalized to the non-treated control (NTC) group. Data are presented as mean ± SEM (N=3, n≥3). **(j)** Confocal fluorescence microscopy analysis of MC38 Gal8-GFP reporter cells incubated with 0.5 μM LDR for 24 h and subsequently irradiated with 532 nm nanosecond pulsed laser at varying fluences: LDR only control (LDR CTR, no laser), 0.3, 0.8, 1.6, and 2.0 J cm^-2^. Representative images show Gal8-GFP puncta formation, indicative of lysosomal membrane permeabilization. GFP channel images were converted to grayscale using ImageJ for enhanced contrast. **(k)** Time-resolved dark-field microscopy images showing sequential frames of HeLa cell spheroids incubated with 1 µM LDR before and after exposure to a 532 nm nanosecond pulsed laser at a fluence of 1.2 J cm^-2^. **(l)** Single time point analysis of spheroids using bright-field microscopy.

We next assessed whether VB formation and collapse translated into effective cell killing. HT1080 cells were incubated with either 0.5 µM or 1 µM LDR for 24 h, followed by irradiation with increasing laser fluences ranging from 0.08 to 1.6 J cm^-2^. Control groups without laser exposure (LDR only) and cells treated with low laser fluences (≤ 0.3 J cm^-2^) did not induce significant cytotoxicity. Starting at 0.5 J cm^-2^, corresponding to the previously identified VB threshold for 1 µM LDR, cell viability dropped substantially, inducing ∼70% cell killing at the highest fluence tested (1.6 J cm^-2^) (Fig. 5g). At LDR concentrations <0.5 µM no significant cell killing was noted (Fig. 5h). Tests performed on HT1080 and CT5.3 cell lines showed consistent trends (Supplementary Fig. 13b,c). Furthermore, we also demonstrated that LDR-assisted cell killing was spatially confined (Supplementary Fig. 13e). Altogether, these findings support a dose- and fluence-dependent cell killing effect mediated by LDR-assisted VB generation.

Following laser irradiation at 1.0 J cm^-2^, in contrast to sunitinib-treated cells, LDR-treated cells exhibited less violent photomechanical effects. Nevertheless, real-time confocal microscopy revealed the emergence of transient intracellular cavities consistent with subcellular VB formation (Fig. 5f). Notably, using MC38-eGFP-galectin8 reporter cells, 1 μM LDR exhibited a progressive increase in galectin puncta formation with mounting laser fluences, indicative of endolysosomal membrane permeabilization (Fig. 5j and Supplementary Fig. 13f). In line with sunitinib, significant caspase activation was only observed at laser fluences above LDR’s VB threshold, reinforcing that VB-induced mechanical damage serves as the primary trigger for subsequent apoptotic cell death (Fig. 5i).

Following 2D cell culture, LDR-induced VB formation was also confirmed in HeLa tumor spheroids (Fig. 5k and Supplementary Movie 8). Given the negligible cytotoxicity of LDR, the spheroid diameter remained unchanged after 24 h incubation with 1 µM LDR (Fig. 5l). Minor morphological changes were detected upon laser treatment, again supporting less violent photomechanical ablation compared to sunitinib. At higher fluences, peripheral cells appeared more dispersed and the spheroid boundary less defined, indicating ablation effects at the spheroid surface. Most importantly, LDR-treated spheroids irradiated with 1.8 J cm^-2^ exhibited a clear loss of calcein-AM signal and disrupted morphology in the laser-exposed region, confirming localized cell death (Supplementary Fig. 13g). Although no macroscopic fragmentation or collapse of the spheroids was observed, a significant reduction in cell viability occurred at fluences above 1.6 J cm^-2^ (Supplementary Fig. 13d), confirming that VB generation in LDR-loaded spheroids can induce substantial cell death.

## Lysosomal VB-mediated tumor ablation in patient-derived tumor models

To further validate the concept of lysosome-mediated VB formation in a complex 3D tumor context, patient-derived neuroblastoma AMC691B-GFP tumoroids were exposed to 1 µM LDR (Fig. 6a). Following a 532 nm wavelength single nanosecond laser pulse, the central region of the tumoroid appeared darker with brightfield microscopy imaging, likely due to extensive light scattering caused by intracellular VBs (Supplementary Fig. 15a). This reduced optical transmission gradually returned to baseline levels within 0.5 seconds, corresponding to VB collapse. These observations were corroborated by dark-field microscopy, clearly showing the emergence of numerous VBs (Fig. 6b and Supplementary Fig. 15b). In line with the data obtained on tumor spheroids, no clear macroscopic disruption of the tumoroid structure was noted. Nevertheless, tumoroid viability was significantly reduced at a laser fluence of 0.9 J cm^-2^ (>VB threshold, Fig. 6c). Notably, upon applying a second laser pulse, neither additional VBs nor further viability reduction could be observed (Supplementary Fig. 15c). This result supports the notion that (1) once photosensitized lysosomes are actuated to generate VBs, they are structurally compromised and (2) LDR-mediated intracellular VB formation and lysosomal disruption is driving cell death induction.

**Fig. 6.**
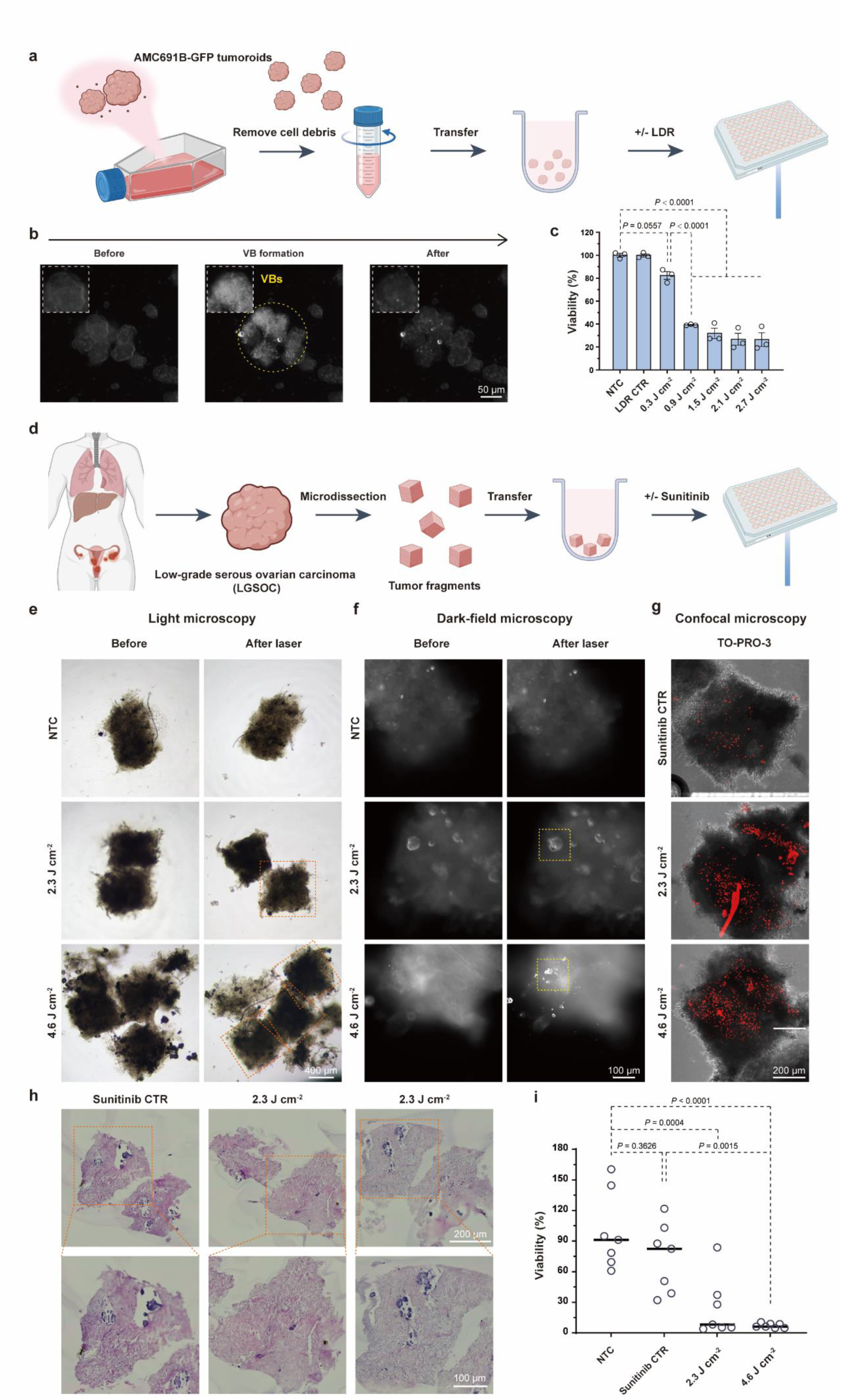
Lysosomal VB-mediated tumor ablation in patient-derived tumoroids and tumor fragments. **(a)** Schematic illustration of the workflow for LDR-treated AMC691B-GFP neuroblastoma tumoroids preparation and culture. **(b)** Dark-field microscopy images of VBs in LDR-incubated tumoroids upon 532 nm laser exposure. **(c)** Quantitative viability analysis of tumoroids treated with varying laser fluences (0.3-2.6 J cm^-2^). Data are presented as mean ± SEM (N=3, n=6). **(d)** Schematic illustration of the workflow for sunitinib-exposed patient-derived tumor fragments preparation and culture. **(e)** Light microscopy images of tumor fragments from a low-grade serous ovarian cancer (LGSOC) patient before and after nanosecond pulsed laser irradiation at fluences of 2.3 and 4.6 J cm^-2^. Non-treated control (NTC) is included for comparison. **(f)** Corresponding dark-field microscopy images revealing the generation of VBs upon laser exposure, with more prominent scattering observed at higher fluence. **(g)** Confocal fluorescence microscopy images of TO-PRO-3-stained tumor fragments showing extent of cell death in the sunitinib only control (sunitinib CTR, no laser) and after irradiation at 2.3 and 4.6 J cm^-2^. **(h)** Representative H&E-stained histological images of tumor fragments. (Top) Untreated tumor fragments incubated with sunitinib only and two groups exposed to a 2.3 J cm^-2^ pulsed laser fluence. (Bottom) Corresponding zoom-in views highlighting local tissue morphology. **(i)** Quantification of tumor fragment’s viability across conditions, confirming increased cell death with mounting laser fluence. Data are presented as median (n=7).

Although such tumoroids capture the three-dimensional cellular architecture of tumors, they still lack the ECM density and heterogeneity of genuine patient-derived tumor tissues, where the dense ECM can hinder both drug diffusion and laser penetration.^57^ Hence, we next investigated ex vivo cultured patient-derived tumor fragments, which preserve the native ECM and cellular composition of clinical tumor specimens. Here, tumor fragments were obtained from a low-grade serous ovarian carcinoma (LGSOC), a tumor type characterized by abundant collagen-rich stroma with small nests of cancer cells. Following a 24 h incubation with 10 μM sunitinib, fragments were irradiated with nanosecond laser pulses at fluences of 2.3 or 4.6 J cm^-2^ (Fig. 6d). Light microscopy imaging revealed increased optical transparency of the tumor fragments post-irradiation compared to the untreated control, suggestive of structural disruption and decreased tumor cell density (Fig. 6e). Real-time dark-field microscopy further confirmed the formation of VBs exclusively in sunitinib-incubated samples. Moreover, VBs were only observed in tumor cell nodules and not in the ECM, further supporting the need for intracellular sunitinib accumulation (Fig. 6f and Supplementary Movie 9 and Movie 10). Confocal microscopy combined with TO-PRO-3 staining revealed markedly higher cell death in the laser-treated group compared to the sunitinib-only control, while preserving the fragment’s structural integrity (Fig. 6g). Consistent with this, H&E staining (Fig. 6h) did not reveal clear signs of structural disruption to the surrounding ECM. Finally, the CellTiter-Glo® 3D cell viability assay quantitatively demonstrated that sunitinib exposure and laser-induced VB generation completely eradicated the tumor cells, even within the densely structured ECM (Fig. 6i). These results collectively confirm that lysosomal accumulation of light-sensitive CADs can be exploited to induce localized tumor ablation, even in clinically relevant, ECM-rich tumor tissues.

## Conclusions

The application of photothermal nanoparticles for cancer tissue ablation often faces several limitations, including poor biodegradability, potential long-term retention in organs such as the liver and spleen, and concerns regarding their immunogenicity and systemic toxicity.^58, 59^ In addition, their limited tumor penetration and uneven intratumoral distribution can compromise treatment uniformity and efficacy.^60^ In this study, we therefore propose a nanoparticle-free strategy for photothermal tumor cell killing by exploiting the lysosomotropic properties of cationic amphiphilic small molecular dyes, as demonstrated for the clinically-approved light-sensitive tyrosine kinase inhibitor (TKI) sunitinib and the commercially available dye Lysotracker™ Deep Red (LDR).^61^

While the lysosomal accumulation of sunitinib has been described to contribute to drug resistance,^36, 37, 62^ it also enables its repurposing as an intracellular photosensitizer. Exposure of a variety of sunitinib-treated cancer cell models to pulsed laser light led to VB formation, depending on the drug concentration and the applied laser fluence. In contrast, laser treatment of free sunitinib in solution (up to 2 mM) does not generate VBs. Based on the pH partition theory for weakly basic drugs, under equilibrium conditions up to 800-fold higher concentrations of sunitinib can be expected in the lysosomal compartment at pH 4.5, compared to the pH neutral cytosol (Supplementary Note 1). It has been demonstrated in literature that CADs can also insert into the membrane of anionic intraluminal vesicles (ILVs) in late endosomal and lysosomal compartments. Moreover, it is known from our earlier work that localized accumulation of dyes may promote their propensity towards VB formation by reducing intermolecular distance. Hence, it is hypothesized that the combination of sunitinib’s optical and lysosomotropic properties converts lysosomes into endogenous photoresponsive substrates for intracellular VB formation, leading to violent photomechanical damage to organelles and/or cells upon their collapse.

Despite a clear correlation between laser application, VB induction and cell killing, other mechanisms cannot be excluded. Both photomechanical, photothermal and photochemical effects can contribute to cell damage evoked by photosensitizers.^63^ In particular, our experiments with sunitinib revealed a certain degree of cell killing at laser fluences below the VB threshold. The latter could be explained by localized production of heat, thus increasing the permeability of the endolysosomal limiting membrane, and therefore promoting cytosolic release of sunitinib from the lysosomes or evoking lysosomal cell death.^10^ Of note, the latter also presents a novel approach to overcoming the challenge of lysosomal entrapment, a major obstacle that may contribute to drug resistance to cationic amphiphilic anticancer drugs. In addition, previous studies demonstrated generation of reactive oxygen species (ROS) upon continuous laser irradiation of intracellular sunitinib.^34, 64^ Nevertheless, our results suggest that VB-mediated cytotoxicity induced by pulsed laser irradiation is not driven by such photochemical mechanisms. Opposed to continuous laser illumination, a single laser pulse delivers energy in a short burst, thus preventing ROS accumulation over time.

Next to sunitinib repurposing, the concept of using lysosomotropic dyes to convert lysosomes into photothermal organelles was additionally validated with the commercial dye LDR, inducing intracellular VBs at a higher wavelength (>500 nm). Our results again demonstrated a clear correlation between intracellular VB formation and cell death, albeit without the extensive photomechanical damage to the treated cells typically observed for sunitinib exposure. Most likely, the subcellular VB-mediated disruption of the lysosomes itself is driving cytotoxicity instead of macroscopic cell ablation. More dedicated experiments would be required to shed light on the exact cell death modalities involved at different experimental conditions.

Finally, we were able to show VB-mediated cell killing with both sunitinib and LDR in tumor spheroids, as well as human patient-derived tumoroids and tumor fragments. Such tumor models more closely resemble the architecture of the tumor microenvironment and provide a more realistic assessment of drug response. Multiple reports have demonstrated limited penetration of nanoparticles of varying sizes in tumor tissue. In contrast, our results reveal that small molecule photosensitizers such as sunitinib and LDR can effectively reach the tumor cells in more complex 3D tumor models and accumulate into their lysosomal compartments, thereby rendering them amenable to VB-mediated cell killing upon pulsed laser application.

To benchmark our nanoparticle-free strategy against conventional photothermal agents, we evaluated the therapeutic efficacy of synthetic nanomaterials in both 2D and 3D HeLa cell models (Supplementary Note 2). Both 80 nm gold nanoparticles (AuNPs) and 150 nm polydopamine nanoparticles (PDNPs) induced higher cytotoxicity in monolayer cultures upon pulsed laser irradiation compared with 3D tumor spheroids, albeit their efficacy remained lower compared to sunitinib. This result highlights the inherent transport barriers and restricted interstitial diffusion of larger synthetic particles within dense tumor architectures, whereas our small-molecule approach ensures homogeneous distribution and effective ablation, even in stroma-rich tumors.

In conclusion, we have identified a nanoparticle-free strategy to enable spatial-selective VB-mediated photoablation of cancer cells using lysosomotropic cationic amphiphilic photosensitizers. As it is based on a universal cellular and physicochemical mechanism, we anticipate that this concept can be utilized across multiple cancer types, photosensitizers and wavelengths, including applications with multiphoton excitation and near infrared (NIR) dyes.

## Materials and methods

### Materials

Dulbecco’s Modified Eagle’s Medium/Nutrient Mixture F-12 (DMEM/F-12), Dulbecco’s Modified Eagle Medium (DMEM, high glucose), Roswell Park Memorial Insitute (RPMI) 1640 medium, Ham’s F-12 Nutrient Mix (Cat. No. 11765054), B-27™ Supplement (50X, serum-free, Cat. No. 17504001), recombinant human FGF-basic (aa 1-155, Cat. No. PHG0266), animal-free recombinant human EGF (PeproTech®, Cat. No. AF-100-15), recombinant human IGF-I (PeproTech®, Cat. No. 100-11), recombinant human PDGF-AA (PeproTech®, Cat. No. 100-13A), recombinant human PDGF-BB (PeproTech®, Cat. No. 100-14B), penicillin/streptomycin solution (100 IU mL^-1^ penicillin, 100 µg mL^-1^ streptomycin), L-glutamine (2 mM) and 0.25% Trypsin-EDTA Gibco™ and Collagen I, rat tail (Cat No. A1048301) were purchased from Gibco®-Life Technologies (New York, USA). Dulbecco’s phosphate-buffered saline without Ca^2+^/Mg^2+^ (DPBS-), bovine serum albumin (BSA) and sodium azide (NaN_3_) were obtained from Sigma-Aldrich (St. Louis, USA). Fetal bovine serum (FBS) was from Hyclone™, GE Healthcare, Machelen, Belgium). Sunitinib malate (CAS No. 341031-54-7) was purchased from Cayman Chemical (Michigan, USA). LysoTracker™ Deep Red (Cat No. L12492), CellROX™ Deep Red (Cat No. C10491), CellROX™ Green (Cat No. C10492) Flow Cytometry Assay Kits, TO-PRO-3 Iodide (Cat No. T3605), Calcein-AM (Cat No. C3099) were obtained from Invitrogen™ (Eugene, USA). The CellTiter-Glo® Luminescent Cell Viability Assay (Cat No. G7571), CellTiter-Glo® 3D Cell Viability Assay (Cat No. G9682) and Caspase-Glo® 3/7 Assay (Cat No. G8091) were obtained from Promega (Belgium).

### Cell lines and culture conditions

Human cervical epithelial adenocarcinoma HeLa cells, lung epithelial A549 cells and T lymphoblast Jurkat E6-1 cells were obtained from American Type Culture Collection (ATCC, Manassas, USA). Fibrosarcoma HT1080 cells and hTERT-immortalized colon cancer-associated fibroblasts (CT5.3hTERT CAFs) were provided by Prof. De Wever (Laboratory of Experimental Cancer Research, Department of Human Structure and Repair, Ghent University). MC38-eGFP-Galectin 8 was provided by Prof. De Geest (Biopharmaceutical Technology Unit, Department of Pharmaceutics, Ghent University). HeLa cells were cultured in DMEM/ F-12, and A549, HT1080 and CT5.3hTERT cells were cultured in DMEM with high glucose. Jurkat E6-1 cells were cultured in RPMI 1640. All culture media were supplemented with 10% FBS, 2 mM L-Glutamine and 100 U mL^-1^ penicillin/streptomycin (*i.e.* complete cell culture medium). All cell lines were maintained in a humidified atmosphere containing 5% CO_2_ at 37°C and culture medium was renewed every other day. Cell lines are regularly tested and found negative for mycoplasma.

### Tumor spheroids

Spheroids were produced via seeding cells in Ultra-low Attachment (ULA) round-bottom 96-well plates (PrimeSurface®, S-BIO, Japan) with a density of 4 × 10^3^ cells per well. The spheroids were incubated in a humidified atmosphere containing 5% CO_2_ at 37°C and culture medium was renewed every day. Spheroid morphology was monitored daily using bright-field microscopy. Acquired images were subjected to automated segmentation and morphometric analysis, including diameter and circularity, using AnaSP software.^52^ On day 3, the spheroids were incubated with sunitinib at concentrations ranging from 0 to 40 μM for 24 h. Following incubation, the spheroids were collected, washed three times with PBS, and transferred for subsequent sample preparation and analysis.

The AMC691B-GFP tumoroids were provided by Prof. Durinck (Pediatric Precision Oncology Lab, Department of Biomolecular Medicine, Ghent University).^65^ Tumoroids were maintained in a specialized tumoroid medium consisting of DMEM (low glucose, GlutaMAX™) and Ham’s F-12 Nutrient Mix (4:1 v/v), supplemented with 2% B-27™ (minus vitamin A), 1% N-2 supplement, 100 U mL^-1^ penicillin, and 100 µg mL^-1^ streptomycin. This medium was further enriched with recombinant human growth factors, including EGF (20 ng mL^-1^), FGF-basic (40 ng mL^-1^), IGF-I (200 ng mL^-1^), PDGF-AA (10 ng mL^-1^), and PDGF-BB (10 ng mL^-1^).

### Preparation and ex vivo treatment of patient-derived tumor fragments

Patient-derived tumor material was applied consistent with our previously reported protocols.^66^ Collection of tissue samples from patients was conducted in accordance with relevant guidelines and approved by the Ethics Committee of Ghent University Hospital (Ref: BC-06978). Patient-derived low-grade serous ovarian carcinoma (LGSOC) tumor fragments were obtained at the time of surgery following informed written consent. Solid tumor lesions were macroscopically selected by a pathologist and immediately placed in ice-cold DMEM LG (cat. no. 31885023, ThermoFisher) supplemented with 10% FBS (Sigma Aldrich), 100 U ml^-1^ penicillin and 100 µg ml^-1^ streptomycin (cat. no. 15070063, Life Technologies). All tumor material was immediately processed by automated cutting into small tumor fragments of 0.125 mm^3^ and placed on ice. After processing, the tumor fragments from different regions within a tumor were mixed to ensure uniform representation of the tumor. Thereafter individual tumor fragments were seeded in pre-warmed supplemented DMEM LG. To prevent evaporation, Ultra-low attachment (ULA) round-bottom 96-well plates (PrimeSurface®, S-BIO, Japan) were sealed with Breathe-Easy semipermeable membranes (cat. no. Z380059, Merck Life Science). Tumor fragments were cultivated for up to 5 days (37°C, 5% CO_2_) under normoxic conditions.

The tumor fragments were first transferred using sterile 200 μL pipette tips into a new Ultra-low attachment (ULA) round-bottom 96-well plates containing pre-warmed culture medium and treated with sunitinib at a final concentration of 10 µM. Following a 24 h incubation, the tumor fragments were carefully transferred to a flat-bottom 96-well plates (VWR®, Avantor, USA). The drug-containing medium was replaced with 200 μL of fresh culture medium per well to remove extracellular sunitinib. Subsequently, the fragments were subjected to laser treatment.

### Quantification of cellular accumulation of sunitinib by flow cytometry

Cells were seeded at a cell density of 8 × 10^3^ cells per well in flat-bottom 96-well plates (VWR®, Avantor, USA) and allowed to adhere overnight. The next day, serial dilutions of sunitinib in cell culture medium were applied at final concentrations ranging from 0 to 10 µM. After incubation for 6, 12 or 24 h at 37°C, lysosomes were labeled by incubation with 75 nM LysoTracker™ Deep Red (LDR) (Thermo Fisher Scientific, Waltham, MA, USA) for 15 min at 37°C. Sunitinib fluorescence, LDR intensity and side scatter (SSC) signal were measured by flow cytometry, using a CytoFLEX flow cytometer equipped with a 96-well plate reader (Beckman Coulter, Krefeld, Germany) and CytExpert software. Data analysis was performed using FlowJo_v10.6.2 (Treestar, Costa Mesa, CA, USA).

### Co-localization of sunitinib with lysosomes observed by confocal microscopy

Cells were seeded in sterilized and tissue culture-treated 96-well microplates (PhenoPlate™, Revvit, The Netherlands) with black well walls and an optically clear cyclic olefin bottom at a density of 1.2 × 10^4^ cells per well and were allowed to adhere overnight. The following day, cells were incubated with different concentrations of sunitinib for 3, 6, 12, 24, 48 or 72 h. Before imaging, sunitinib-containing cell culture medium was removed and cells were stained with 75 nM Lysotracker™ Deep Red (LDR) for 15 min at 37°C and the nuclei were labeled with Hoechst 33342 (Molecular Probes, Belgium) in cell culture medium (1/2000 dilution) for 15 min at 37°C. Microscopy imaging was performed with a laser scanning confocal microscope (Nikon A1R HD confocal, Nikon, Japan), equipped with a 60× water objective lens (60× WI Plan Apo, NA 1.27, WD 180 μm, Nikon, Japan) with a laser box (LU-N4 LASER UNIT 405/488/561/640, Nikon Benelux, Brussels Belgium) and detector box (A1-DUG-2 GaAsP Multi Detector Unit, GaAsp PMT for 488 and 561 and Multi-Alkali PMT for 640 and 405 nm). The 405 nm, 488 nm and 561 nm laser were applied to excite the Hoechst labeled nuclei, sunitinib and LDR, respectively. Fluorescence emission was detected through 450/50 nm (MHE57010), 525/50 nm (MHE57030) and 595/50 nm (MHE57050) filter cubes, respectively. A Galvano scanner was used for unidirectional scanning to acquire the sequential channels with 2 times line averaging and scan speed of 0.042 FPS. The pinhole was set to 20.43 µm and the pixel size was 100 nm pixel^-1^. NIS Elements software (Nikon, Japan) was applied for imaging. Images were analyzed with ImageJ (FIJI, https://fiji.sc/).^67^

### Laser irradiation and vapor bubble formation

Vapor bubble (VB) detection was performed using a previously reported in-house developed setup including an optical system and an electric timing system.^56, 68^ The system includes a pulsed laser source (Opolette HE 355 LD, OPOTEK Inc., CA, USA) with a pulse duration of approximately 7 ns, a repetition rate of 20 Hz, and a beam diameter of 222 µm. The excitation wavelength was set to 480 nm. Cells were seeded at a density of 1 × 10^5^ cells per dish onto 50 mm glass-bottom Petri dishes (MatTek, P50G-1.5-30-F) that had been pre-coated with Gibco™ Collagen I, rat tail (Thermo Fisher Scientific) for 1 h at room temperature. After 24 h of incubation to allow cell attachment, cells were treated with varying concentrations of sunitinib (0-20 µM) and further incubated for an additional 24 h prior to laser irradiation. VBs were visualized via dark-field microscopy using a dark-field condenser, capitalizing on the strong light scattering properties of nano/micro bubbles. An EMCCD camera (Cascade II: 512, Photometrics, Tucson, AZ, USA) was synchronized with the laser pulses through an electronic pulse generator (BNC575, Berkeley Nucleonics Corporation, CA, USA), enabling the acquisition of images before, during, and after laser exposure. Laser pulse energy was monitored in real time using an energy meter (J-25MB-HE&LE, EnergyMax-USB/RS sensors, Coherent), and laser fluence was calculated as the average energy per pulse divided by the beam area (J cm^-2^). By counting the number of VBs within the laser irradiation area for increasing laser fluence, one can determine the VB threshold, defined as the laser pulse fluence at which 90% of the plateau of producing VB.^69^ To precisely determine this threshold and characterize the VB generation dynamics, the experimental data were fitted using a Boltzmann sigmoidal model:

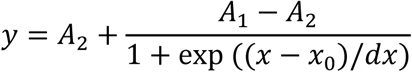

In this equation, *A*_0_ and *A*_1_represent the lower and upper plateaus of the VB count, respectively, while *x*_0_ denotes the inflection point (midpoint) of the transition and *d*_*x*_ signifies the width of the response curve. The VB generation threshold *x*_90_ was then calculated as *x*_90_ = *x*_0_ + *d*_*x*_ · ln(9), representing the laser fluence required to reach 90% of the saturation plateau. Furthermore, based on these fitting parameters, the VB induction process was categorized into three distinct regimes: (i) the “Sparse” regime, spanning from the baseline *A*_1_to the inflection point *x*_0_, characterized by a low and stochastic VB count; (ii) the “Moderate” regime, defined from *x*_0_ to the *x*_90_ threshold, where the VB number increases significantly and non-linearly; and (iii) the “Threshold” regime (above *x*_90_), where frequent and stable VB formation occurs, leading to consistent photomechanical effects.

To facilitate the identification and quantification of individual VBs against the background, frame-by-frame image analysis was performed using ImageJ (FIJI, https://fiji.sc/), allowing precise visualization of VB formation and collapse dynamics. In addition to EMCCD-based detection, an sCMOS camera was also used for real-time monitoring of VB formation and threshold measurements. To support reproducibility and visualization, the operation window of the Micro-Manager software was screen-recorded using Free Cam (Digital Thoughts Software), and representative videos have been provided in the Supplementary Videos.

### Programmatic area-defined laser scanning

For quantification of cell viability as a function of laser irradiation, HeLa cells were seeded at a cell density of 8 × 10^3^ cells per well in flat-bottom 96-well plates (VWR®, Avantor, USA) and were allowed to adhere overnight. The following day, sunitinib dilutions in cell culture medium at final concentrations ranging from 0 to 10 µM were applied, and cells were incubated for 24 h at 37°C. For laser treatment, the plates were placed on an electronic motorized translation stage (H117, Prior, UK), enabling automated laser scanning across each well. The laser was scanned in a "snake"-type pattern (Supplementary Fig. 9a), with a scanning speed of 7 × 10^3^ µm s^-1^. The beam was moved line by line across the well with a fixed step width of 0.2 mm in the Y-direction, matching the laser spot diameter to ensure continuous, non-overlapping coverage of the entire well. X-field and Y-field parameters were user-defined to control the number of scans in each direction, and the edge of the 96-well plate was used as a reference point to initialize each scanning sequence. This setup ensured that the laser beam uniformly covered the entire well surface without redundancy or omission.

### Evaluation of cellular health

Cell viability was then measured via detection of cell’s metabolic activity using a CellTiter-Glo® assay (Promega, Belgium) following the manufacturer’s instructions. Correspondingly, the viability of spheroids was measured using the CellTiter-Glo® 3D Cell Viability Assay. Data are presented as a percentage of viable cells calculated from the luminescence signal of each condition relatively to non-treated cells and by taking into account the background fluorescence of the medium.

Apoptosis was assessed via detection of cell’s caspase activity using the Caspase-Glo® 3/7 assay (Promega, Belgium) following the manufacturer’s instructions. Briefly, cells were seeded in 96-well plates (VWR®, Avantor, USA) and incubated under standard conditions. Prior to laser irradiation, the culture medium containing sunitinib was removed and replaced with 100 μL of fresh culture medium. Immediately after laser treatment, the plates were returned to the incubator and cells were allowed to recover for an additional 6 h. Subsequently, 100 μL of Caspase-Glo® 3/7 reagent was added directly to each well, bringing the total volume to 200 μL. Plates were gently shaken at 100 rpm for 10 min and then incubated at room temperature for 20 min to allow luminescence development. Following incubation, 150 μL of the supernatant was transferred from each well into a new white-walled 96-well microplate (PS, F-Bottom, Chimney Well, Greiner, Germany), and luminescence was measured using a GloMax® Microplate Reader. Caspase activity was expressed as relative caspase-3/7 activity (fold of control).

### Confocal microscopy imaging of spatial-selective cell killing

HeLa cells and A549 cells were irradiated according to a pre-defined pattern. Two vertical lines with a width of 1 mm, separated by a non-irradiated zone of 100 µm (Supplementary Fig. 9b), were scanned and cells were subsequently incubated for 4 h at 37°C, 5% CO_2_. After incubation, cells were stained with 0.5 µg mL^-1^ calcein-AM green cell-permeant dye (Invitrogen) and 1 µM TO-PRO-3 Iodide (Invitrogen) nucleic acid stain for 15 min at 37°C, 5% CO_2_, to identify viable and necrotic cells respectively, and afterwards visualized using confocal microscopy.

Widefield fluorescence and spinning disk confocal images were recorded with a Nikon Ti microscope using an Andor DU-897 X-8026 camera with a 20× objective lens (Plan Apo λ 20× 0.45NA, Nikon), numerical aperture of 0.75 and a refractive index of 1. Two planes were imaged at 488/640 nm. The camera settings included a 1x1 binning, 100 ms exposure, and a multiplier of 300. The readout speed was set to 17 MHz with a conversion gain of Gain 1, and the vertical shift speed was 3.3 µs. The PFS-S system was active with an offset of 8094 and the mirror inserted. The illumination was controlled via the NIDAQ system. Imaging was performed at a scan speed of 4900 rpm using the CSU system, with the CSU shutter open and fiber output set to EPI. Image acquisition was carried out in multi-excitation mode, ensuring precise control of the laser lines and their corresponding wavelengths. NIS Elements was used as acquisition software. A large field-of-view image was composed of 9 by 9 images (512 × 512 image size) with 15% overlapping and stitching by a blending algorithm. Images analysis was performed using ImageJ (FIJI, https://fiji.sc/), including correction of field non-uniformity.

### Reactive oxygen species detection

Intracellular reactive oxygen species (ROS) generation was assessed using the CellROX™ Deep Red Reagent (Thermo Fisher Scientific), a cell-permeant fluorogenic probe that becomes highly fluorescent upon oxidation by ROS. The dye exhibits excitation/emission maxima at approximately 640/665 nm, allowing for reliable detection of oxidative stress in live cells via flow cytometry and confocal fluorescence microscopy.

To avoid interference from trypsinization-induced oxidative stress, suspension Jurkat cells were used. Cells were first incubated with either sunitinib or LysoTracker™ Deep Red (LDR) at indicated concentrations for 24 h. After incubation, cells were pelleted by centrifugation, washed once with PBS, and resuspended in fresh medium containing 5 µM CellROX™ Deep Red. Laser treatment was immediately applied following dye addition. Subsequently, 1 mM SYTOX® Blue Dead Cell Stain (Thermo Fisher Scientific) was added to each sample and incubated for 30 min at 37°C to distinguish live and dead cells.

Samples were then analyzed by flow cytometry (CytoFLEX, Beckman Coulter) or imaged by confocal microscopy. ROS generation was evaluated based on the intensity of deep red fluorescence in CellROX™-positive cells, while SYTOX® Blue was used concurrently to assess membrane integrity. No-laser control was included to determine baseline ROS levels in the absence of photothermal stimulation.

### Transmission electron microscopy (TEM) sample preparation and imaging

HeLa cells were cultured on acid-cleaned glass coverslips (BRAND®, 20 mm × 20 mm square) for TEM analysis. Coverslips were pre-treated by overnight immersion in concentrated sulfuric acid (ACS reagent, 95.0-98.0%), followed by extensive rinsing with RNase-free water and thorough drying. To ensure stable adhesion of the coverslip during cell seeding, 10 μL of PBS was first applied to the center of a 12-well plate (VWR®, Avantor, USA), and the coverslip was placed onto the droplet. This improved surface wetting and prevented displacement during medium addition. A 100 μL cell suspension containing 4 × 10^4^ HeLa cells was then gently dispensed onto the coverslip. After a 30 min settling period to allow cell sedimentation onto the coverslip, 1 mL of fresh culture medium was slowly added along the inner wall of the well. Cells were incubated for 24 h before treatment. Following initial attachment, the culture medium was replaced with medium containing 10 μM sunitinib for 24 h. After incubation, the medium was removed and replaced with fresh medium prior to laser treatment. Using a microscope integrated with the nanosecond pulsed laser system, the coverslip region was identified and subsequently irradiated at specified fluences (0.3 or 0.5 J cm^-2^). The cells on coverslips were fixed using a two-step primary fixation protocol. First, culture medium was gently replaced with a 1:1 mixture of fresh medium and primary fixative solution (4% formaldehyde [EM-grade], 2.5% glutaraldehyde [EM-grade] in 0.1 M cacodylate buffer), followed by gentle shaking for 10 min at room temperature. This was then replaced by 100% fixative solution, and cells were further fixed for 1 h at room temperature. After fixation, samples were washed three times for 10 min each at 4°C with 0.1 M cacodylate buffer. Post-fixation was performed in reduced 1% osmium tetroxide solution (0.132 g K_3_[Fe(CN)_6_] + 2 mL 4% OsO_4_ + 6 mL 0.1 M cacodylate buffer) for 1 h on ice with gentle shaking. Samples were subsequently washed three times in distilled water (3 × 10 min, 4°C, shaking), then stained *en bloc* in 1% uranyl acetate at 4°C under light-protected conditions. Dehydration was carried out at room temperature with sequential ethanol solutions (15%, 30%, 50%, 70%, 95%, and two changes of 100% ethanol), each for 15 min with gentle shaking. Resin infiltration was performed at room temperature using Spurr’s resin mixed with 100% ethanol in increasing ratios (1:3, 2:3, and 3 × 3:3), with each step lasting at least 8 h under continuous shaking. For embedding, coverslips were gently blotted to remove excess resin and inverted onto freshly filled embedding molds containing Spurr, with the cell side facing downward. Samples were polymerized at 69°C for approximately 16 h. Ultrathin sections (∼70 nm) were obtained using a Leica EM UC6 ultramicrotome. Subsequently, the sections were post-stained with 1% uranyl acetate for 40 min and 3% lead citrate for 10 min using a Leica EM AC20 automatic stainer. Images were acquired using a JEOL JEM-1400Plus transmission electron microscope operating at an acceleration voltage of 80 kV. Morphological changes were assessed with a focus on lysosomal structure and subcellular damage induced by laser treatment. Lysosome diameters were quantified across different treatment groups.

### Statistics and reproducibility

Experiments were performed in biological triplicate (N = 3), each including at least three technical replicates, and results are presented as mean ± standard error of the mean (SEM), unless stated otherwise. Statistical analysis was performed using the 9th version of the GraphPad Prism software. One-way ANOVA with Tukey Correction was applied to compare multiple conditions.

## Data availability

All the data that support the findings of this study are available within the Article and its Supplementary Information. Source data are provided with this paper.

## Acknowledgements

The authors gratefully acknowledge the Ghent Light Microscopy Core (GLiM), the VIB HistoCore, and the VIB BioImaging Core (VIB-UGent Center for Inflammation Research) for providing access to their facilities and technical support. This research was supported by the China Scholarship Council (Grant No. 202208320061), the European Research Council (ERC) under the European Union’s Horizon Europe research and innovation program (Grant Agreement No. 101075873, DYE-LIGHT), and the EIC Pathfinder Grant NOVISTEM (Grant Agreement No. 101071105). Furthermore, the authors acknowledge the financial contributions from the Foundation against Cancer (Stichting tegen Kanker), Kom op Tegen Kanker (Grant No. STI.VLK.2022.0003.01), and Villa Joep (Grant No. STI.DIV.2024.0003.01).

## Author contributions

T.L., C.M., S.D.S., K. Remaut and K. Raemdonck conceived the idea, formulated research goals and planned the experiments. T.L. and C.M. performed most of the experiments and data analysis and wrote the manuscript. D.P. provided guidance on the setup, calibration, and confocal microscopy related to the pulsed laser system. F.S. and K.B. provided guidance and support for evaluating laser-induced cell killing and photoablation in tumor spheroids. H.D.K. provided technical support and guidance for confocal fluorescence microscopy. R.D.R., F.B., and K.L. assisted in TEM sample preparation, image acquisition, and histological analysis. C.D.C. and W.L. prepared the spheroid samples and assisted in experimental procedures. B.V. provided support and advice on ROS detection methods. O.D.W. and K.D. contributed to the design and interpretation of tumor model experiments. O.D.W. and P.T. provided patient-derived tumor fragments and the associated clinical expertise essential for this study. K. Remaut, S.D.S, K.B. and K. Raemdonck supervised the research, advised on overall data analysis and finalized the manuscript. All authors discussed the results and contributed to the final version of the manuscript.

