## Supplementary Information for "Converting Lysosomes into Photothermal Organelles Enables Nanoparticle-Free Tumor Ablation via Intracellular Vapor Bubbles"

### Supplementary Notes

#### Supplementary Note 1 Sunitinib pH Partition Theory Calculation

**Parameters:**

- cLogP: 5.2
- pKa: 9.0
- Lysosomal pH: 4.5
- Cytosolic pH: 7.4

**Calculation Steps:**

According to the pH partition theory, under equilibrium conditions during cell exposure to sunitinib, the following equation can be written for a weakly basic drug:

$$\frac{C_{lysosome}}{C_{cytosol}}=\frac{1+antilog({pK}_{a}-4.5)}{1+antilog({pK}_{a}-7.4)}$$

Given the abovementioned parameters,

$$\frac{C_{lysosome}}{C_{cytosol}}\approx775$$

This means that sunitinib is expected to accumulate approximately 800 times more in the lysosomes at pH 4.5 compared to the cytosol or extracellular medium at pH 7.4.

### Supplementary Note 2

**Photothermal therapy with synthetic nanoparticles in 2D and 3D models**

HeLa cells were seeded in 96-well flat-bottom plates at a density of 1.2 × 10^4^ cells per well for 2D monolayer cultures, and in Ultra-low Attachment (ULA) round-bottom 96-well plates (PrimeSurface®, S-BIO, Japan) at 4 × 10^3^ cells per well to promote spheroid formation. After 24 h of incubation, reaching approximately 90% confluency for monolayers and a diameter of ~200 µm for spheroids, 100 µL of the culture medium was replaced with nanoparticle solutions of varying concentrations to achieve final concentrations of 10^6^, 10^7^, 10^8^, and 10^9^ NPs mL^-1^. Citrate-stabilized gold nanoparticles (citrate-AuNPs, 80 nm, Lot No. CMAU802534000001-01) and BSA-coated polydopamine nanoparticles (PDNPs-BSA, 150 nm, Lot No. 2405001-pd-150) were purchased from Trince Bio (Ghent, Belgium). Following a 24 h incubation period, non-internalized nanoparticles were removed by washing the cells three times with fresh culture medium. For laser treatment using the Lumipore™ device (Trince, Ghent, Belgium), spheroids were carefully transferred from the U-bottom plates to flat-bottom 96-well plates (VWR®, Avantor, USA) using a 200 µL pipette. Both 2D and 3D cultures were then subjected to pulsed laser irradiation at a fluence of 2.0 J cm^-2^. After irradiation, the plates were incubated for an additional 4 h at 37 °C. Finally, cell viability was quantified respectively using the CellTiter-Glo® and CellTiter-Glo® 3D cell viability assays (Promega, Belgium) according to the manufacturer’s instructions.

**Supplementary Movies**

**Supplementary Movie 1. VBs_HeLa_10 μM Sunitinib_0.8 J cm^-2^**

This movie shows the generation and collapse of vapor bubbles (VBs) in HeLa cells incubated with 10 µM sunitinib for 24 h. A single laser pulse at 0.8 J cm^-2^ was applied, capturing the rapid bubble formation and subsequent collapse events at the subcellular level.

**Supplementary Movie 2. 2 mM Sunitinib Control_pH 5_2.0 J cm^-2^**

This movie shows a control experiment in which 2 mM sunitinib was dissolved in pH 5 acetate buffer and imaged under dark-field microscopy. A single laser pulse at 2.0 J cm^-2^ was applied to visualize the photothermal response of freely dissolved sunitinib in the absence of cells.

**Supplementary Movie 3. VBs_Jurkat_10 μM Sunitinib_0.8 J cm^-2^**

This movie shows the generation and collapse of VBs in Jurkat cells incubated with 10 µM sunitinib for 24 h. A single laser pulse at 0.8 J cm^-2^ was applied, capturing the rapid photothermal bubble formation and subsequent collapse in suspension cells.

**Supplementary Movie 4. VBs_HeLa_5 μM Sunitinib_0.8 J cm^-2^**

This movie shows VB formation and collapse in HeLa cells incubated with 5 µM sunitinib for 24 h. A single laser pulse at 0.8 J cm^-2^ was applied, allowing visualization of subcellular bubble generation and collapse at a lower intracellular drug loading compared to the 10 µM condition.

**Supplementary Movie 5. VBs_HeLa_20 μM Sunitinib_0.8 J cm^-2^**

This movie shows the formation and collapse of VBs in HeLa cells incubated with 20 µM sunitinib for 24 h. A single laser pulse at 0.8 J cm^-2^ was applied, illustrating the photothermal bubble dynamics at a higher intracellular drug accumulation.

**Supplementary Movie 6. VBs_HeLa Spheroids_10 μM Sunitinib_1.0 J cm^-2^**

This movie shows VB formation in a HeLa spheroid incubated with 10 µM sunitinib for 24 h. A single laser pulse at 1.0 J cm^-2^ was applied, visualizing localized VB generation within the 3D tumor spheroid structure.

**Supplementary Movie 7. VBs_HT1080_1 μM LDR_1.2 J cm^-2^**

This movie shows VB formation in HT1080 cells incubated with 1 µM Lysotracker™ Deep Red (LDR) for 24 h. A single laser pulse at 1.2 J cm^-2^ was applied to visualize the photothermal response of LDR-loaded lysosomes in this cancer cell line.

**Supplementary Movie 8. VBs_HeLa Spheroid_1 μM LDR_532nm_1.2 J cm^-2^**

This movie shows VB formation in a HeLa spheroid incubated with 1 µM Lysotracker™ Deep Red (LDR) for 24 h. A single 532 nm laser pulse at 1.2 J cm^-2^ was applied, capturing localized VB generation within the 3D spheroid following lysosomal accumulation of LDR.

**Supplementary Movie 9. Tumor fragments_No sunitinib_2.3 J cm^-2^**

This movie shows laser irradiation of tumor fragments (low-grade serous ovarian cancer) without sunitinib incubation. A single laser pulse was applied under the same conditions used for treated samples, demonstrating the absence of VBs formation or detectable photothermal response in untreated tissue.

**Supplementary Movie 10. Tumor fragments_10 μM Sunitinib_2.3 J cm^-2^**

This movie shows tumor fragments incubated with 10 µM sunitinib for 24 h and subsequently irradiated with nanosecond laser pulses. It captures VB formation within the tumor nodules and the resulting localized photomechanical ablation.

**Supplementary Figures**

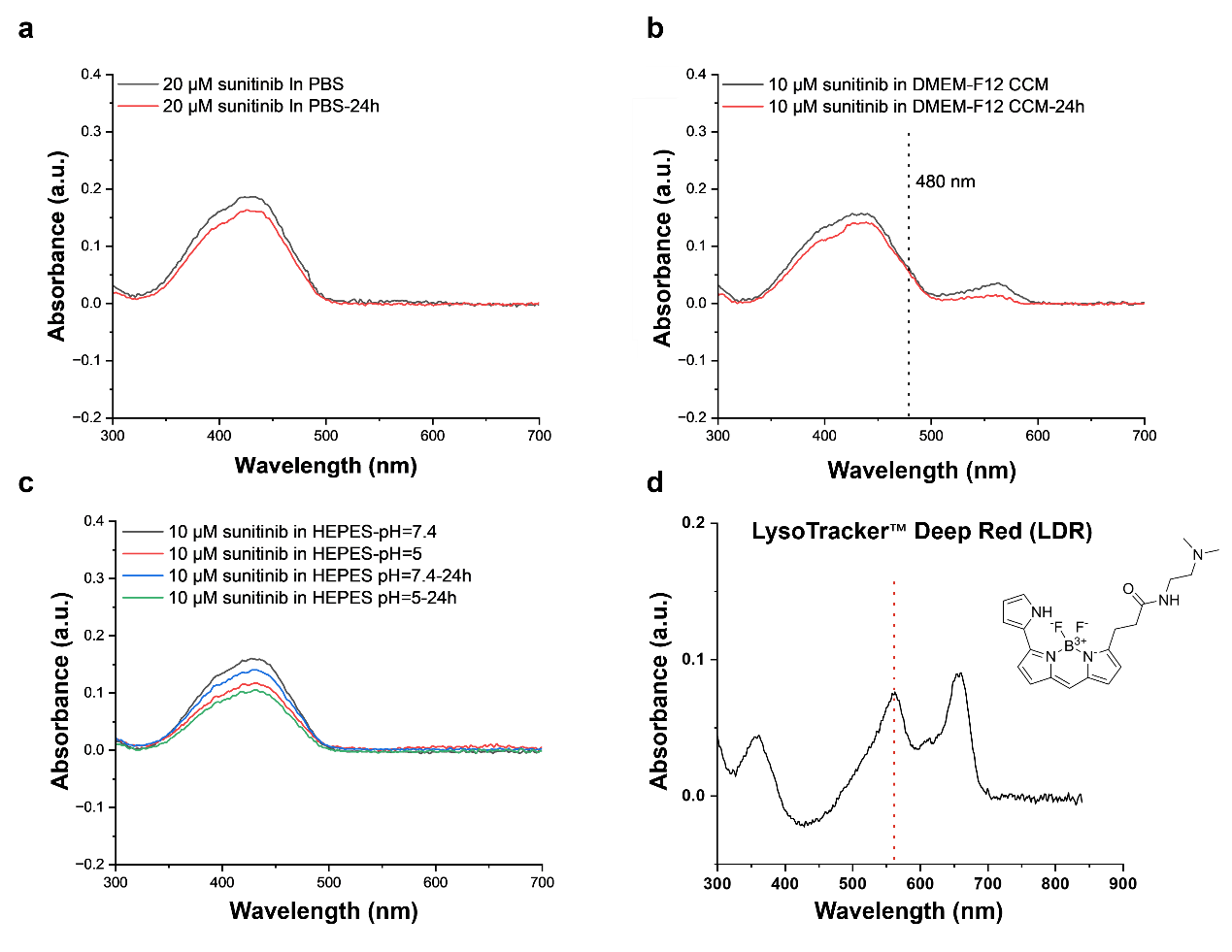

**Supplementary Fig. 1** UV-Vis spectra of sunitinib (malate) in **(a)** PBS, **(b)** DMEM-F12 cell culture medium (CCM) and **(c)** HEPES buffers with varying pH. **(d)** Chemical structure of LysoTracker™ Deep Red and UV-Vis spectra in DMEM-F12 cell culture medium.

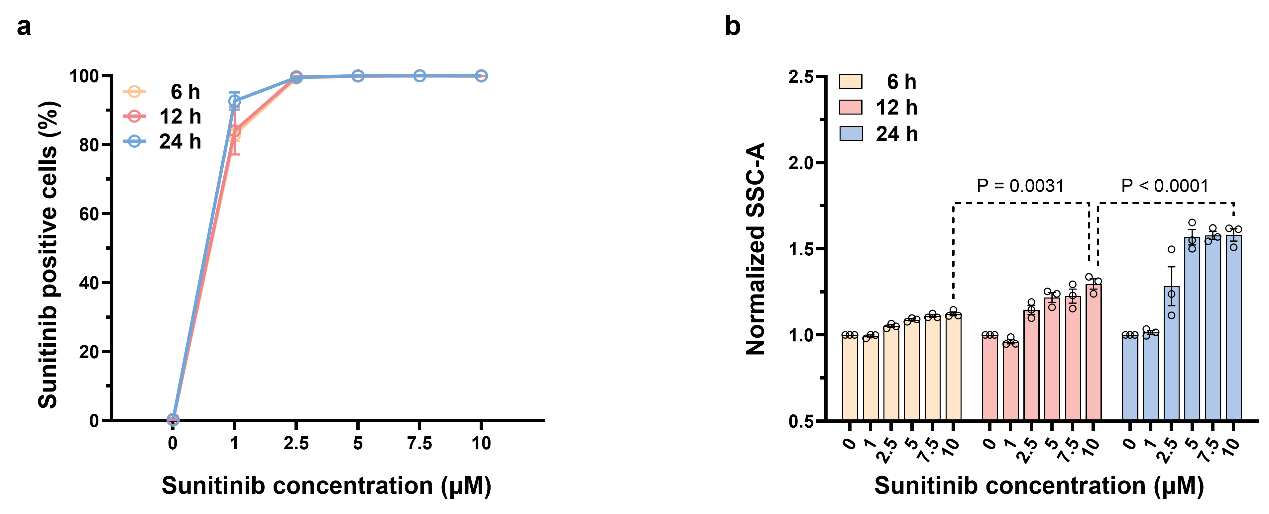

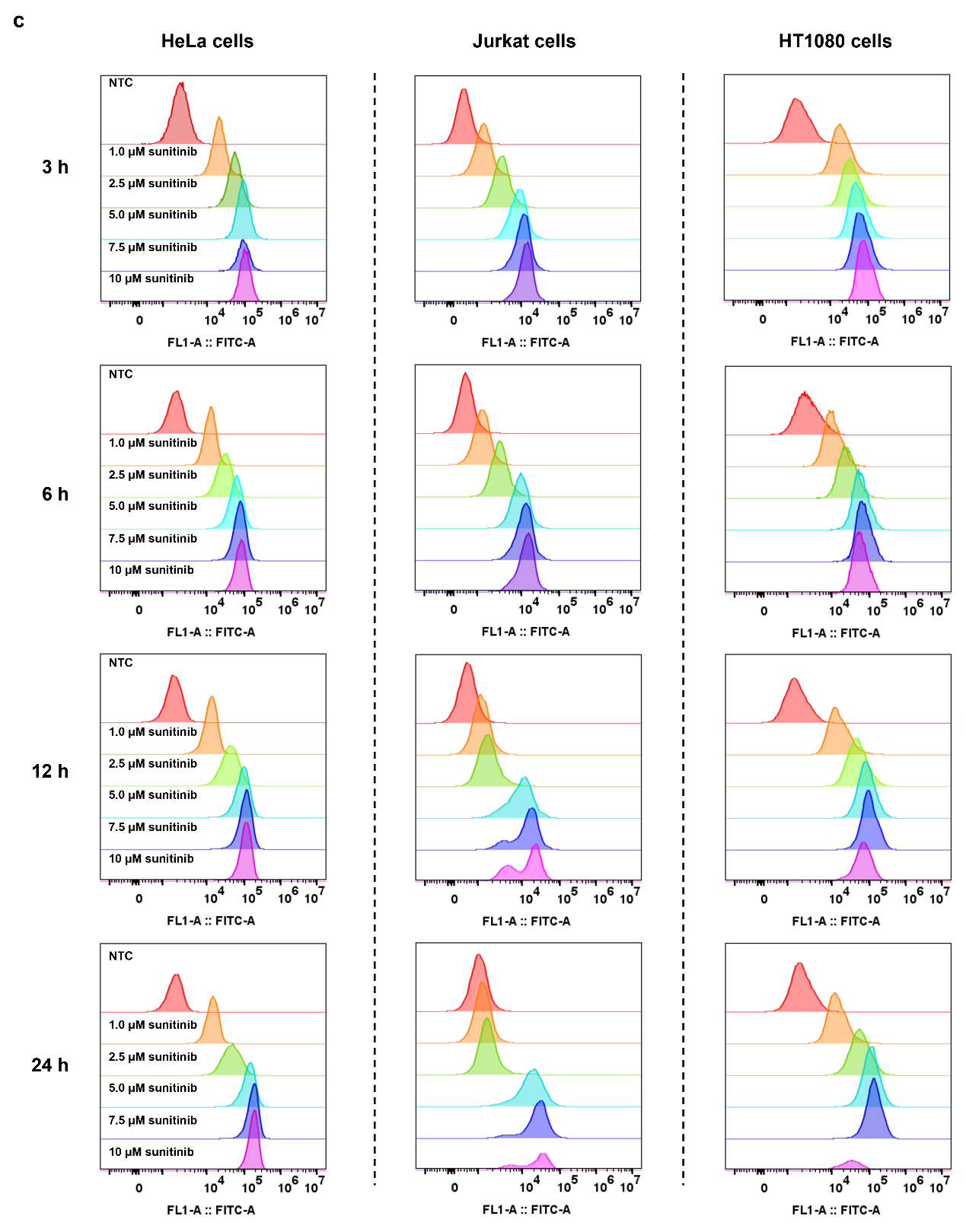

**Supplementary Fig. 2** **(a)** Percentage of sunitinib-positive HeLa cells at different sunitinib concentrations (0-10 µM) and incubation times (6 h, 12 h, 24 h). **(b)** Relative increase of side scatter signal (SSC-A) of HeLa cells as a function of sunitinib concentration and incubation time. Data are presented as mean ± SEM (N=3, n=3). **(c)** Overlaying histograms of the time-dependent and concentration-dependent intracellular fluorescence intensity of sunitinib in HeLa cells, Jurkat cells and HT1080 cells.

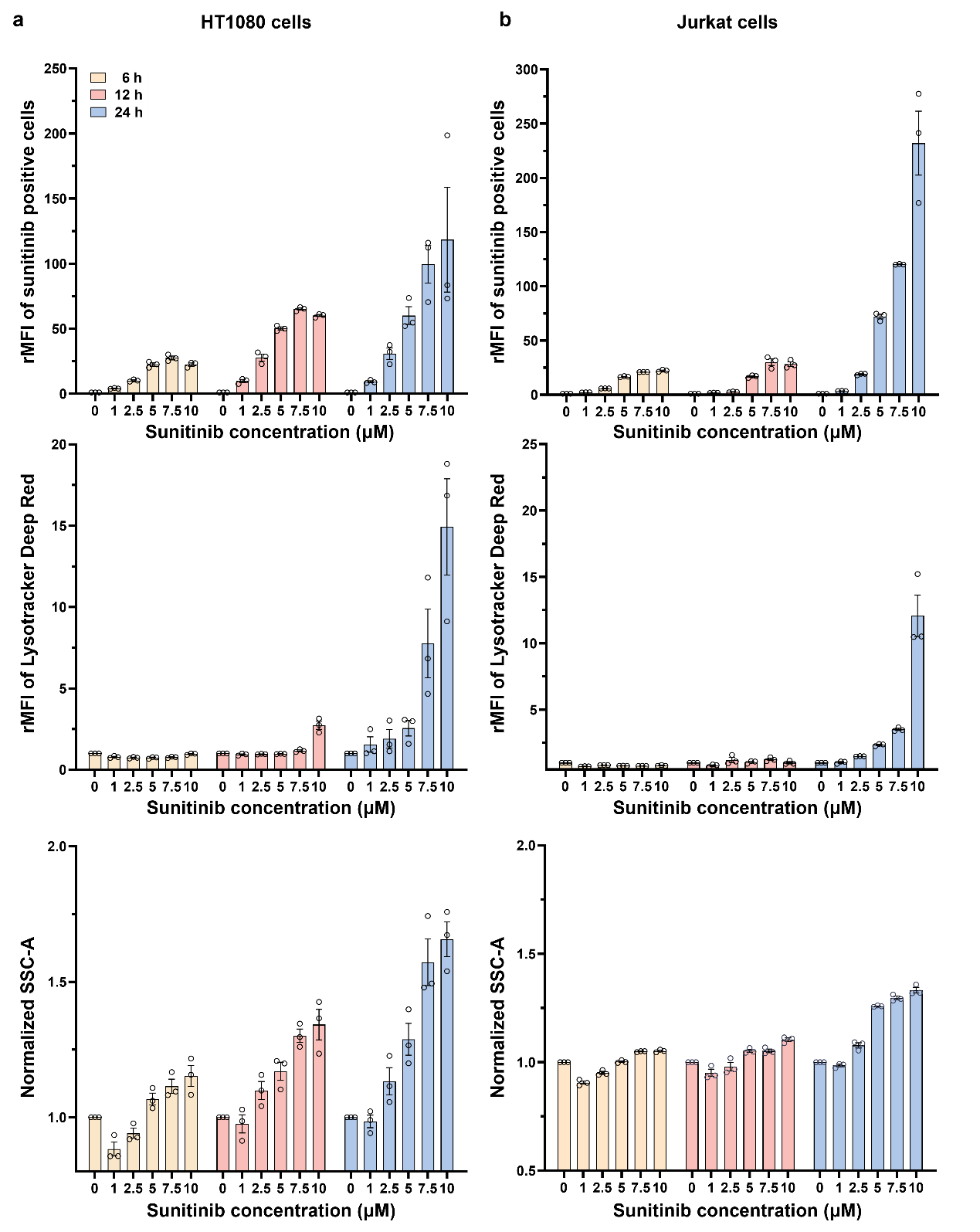

**Supplementary Fig. 3** Characterization of sunitinib accumulation in **(a)** HT1080 cells and **(b)** Jurkat cells. Relative mean fluorescence intensity (rMFI) of cells accumulating sunitinib. Relative increase in Lysotracker™ Deep Red (LDR) staining of cells as a function of sunitinib concentration and incubation time. Relative increase in side scatter signal (SSC-A) of cells as a function of sunitinib concentration and incubation time. Data are presented as mean ± SEM (N=3, n=3).

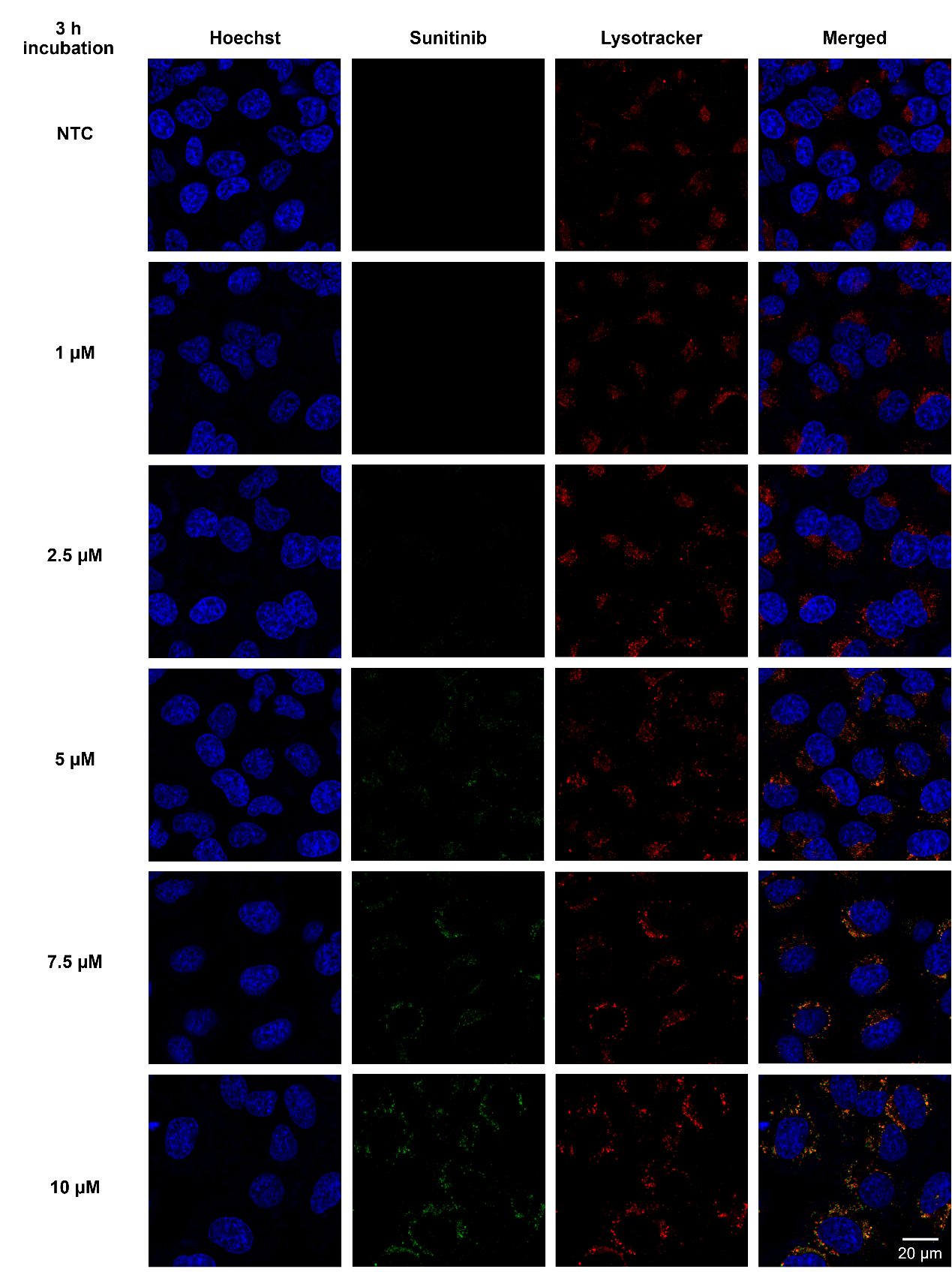

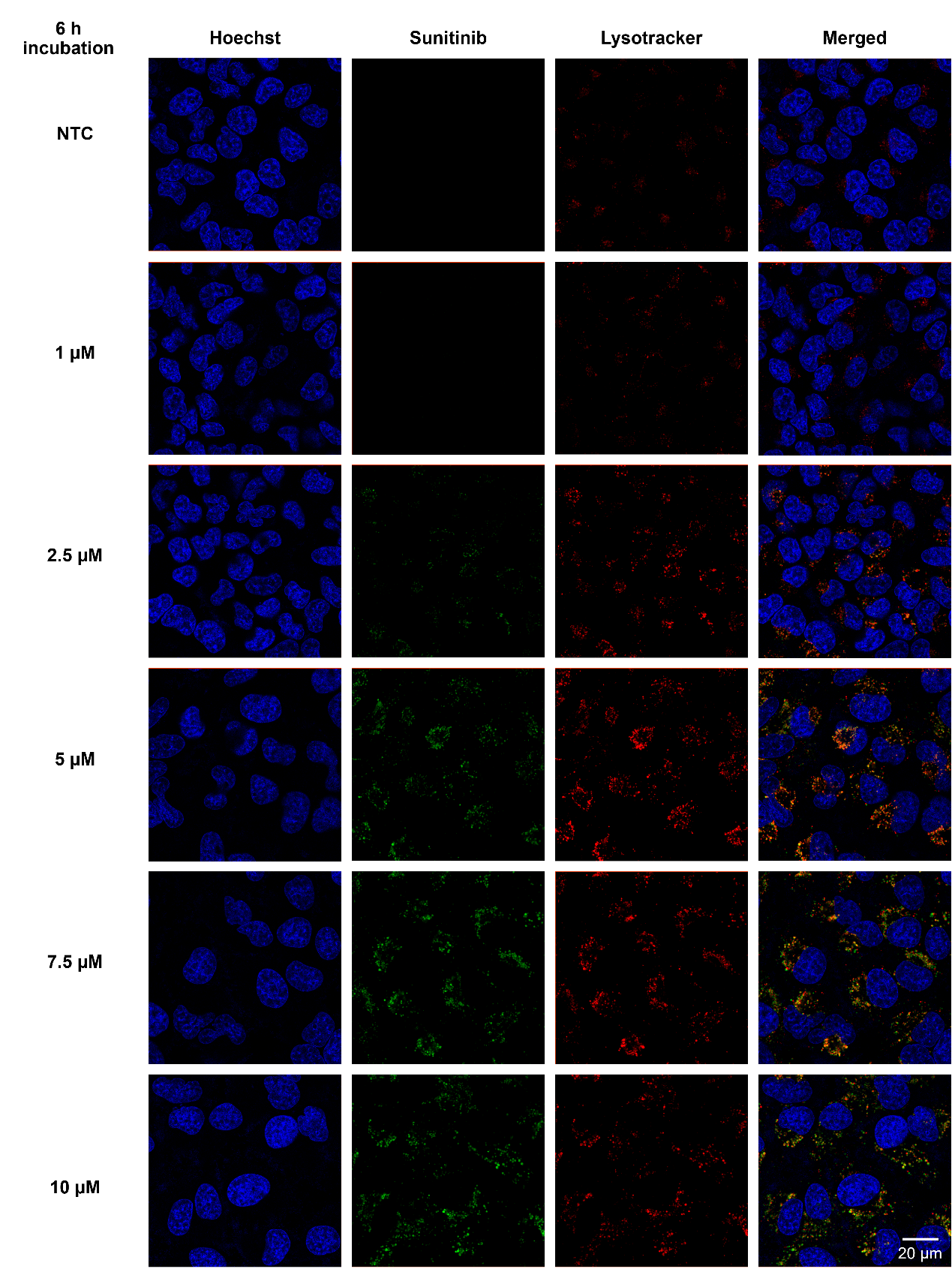

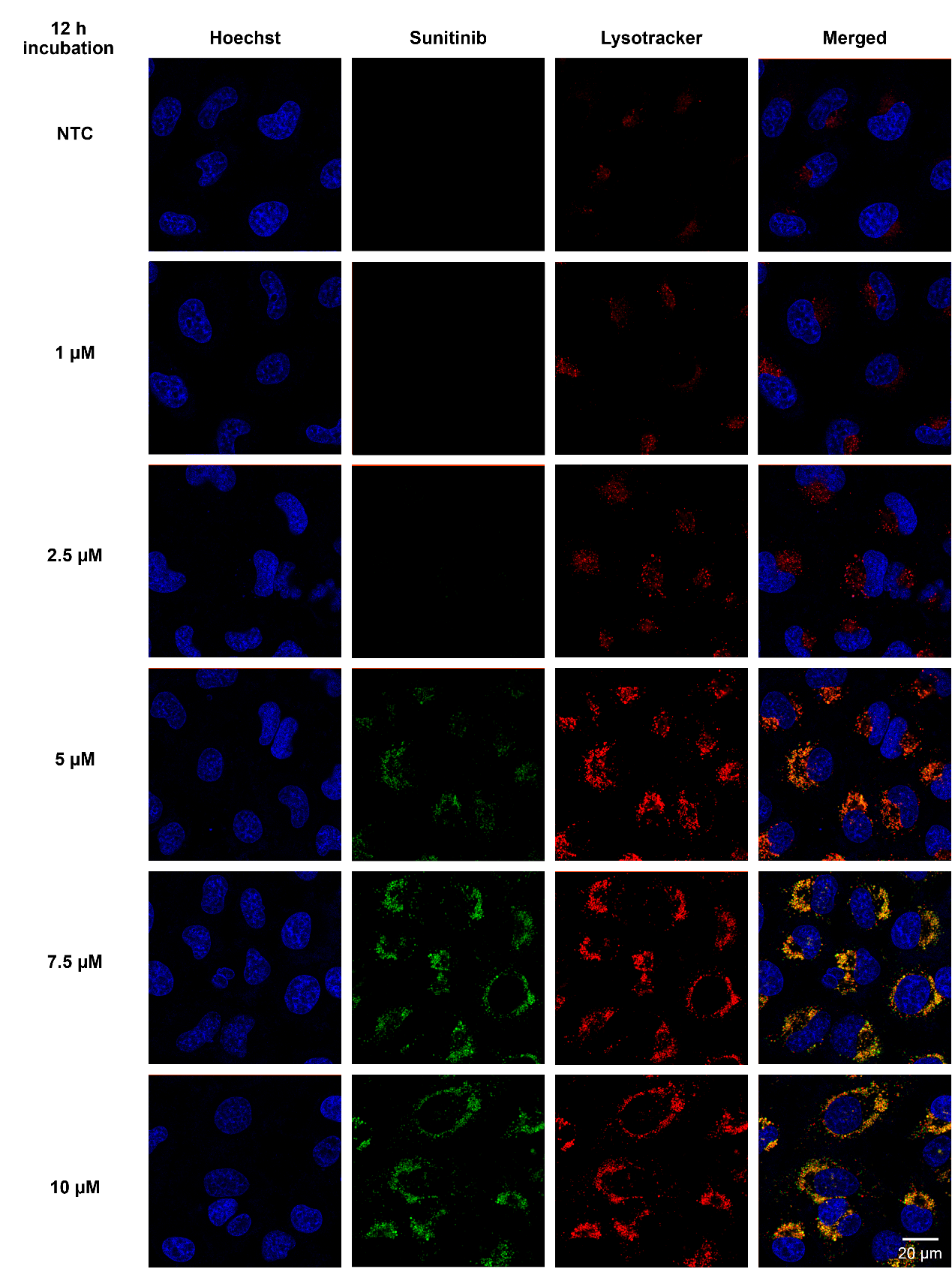

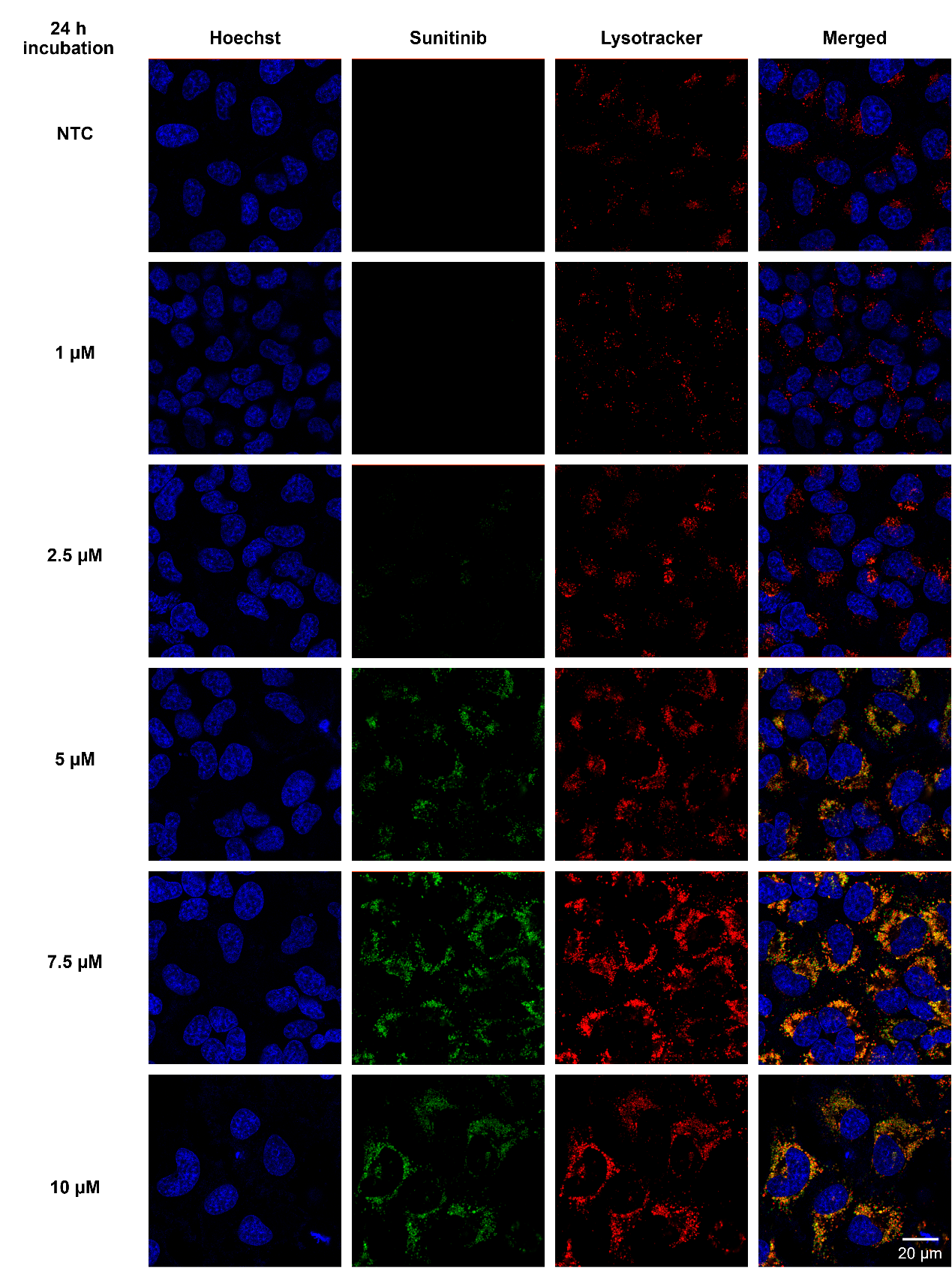

**Supplementary Fig.** **4 Sunitinib accumulation and co-localization with Lysotracker™ Deep Red staining.** HeLa cells were incubated for 3 h, 6 h, 12 h and 24 h with varying concentrations of sunitinib (1 to 10 µM) and subsequently imaged with confocal microscopy following nuclear staining with Hoechst and lysosomal staining with Lysotracker™ Deep Red (LDR).

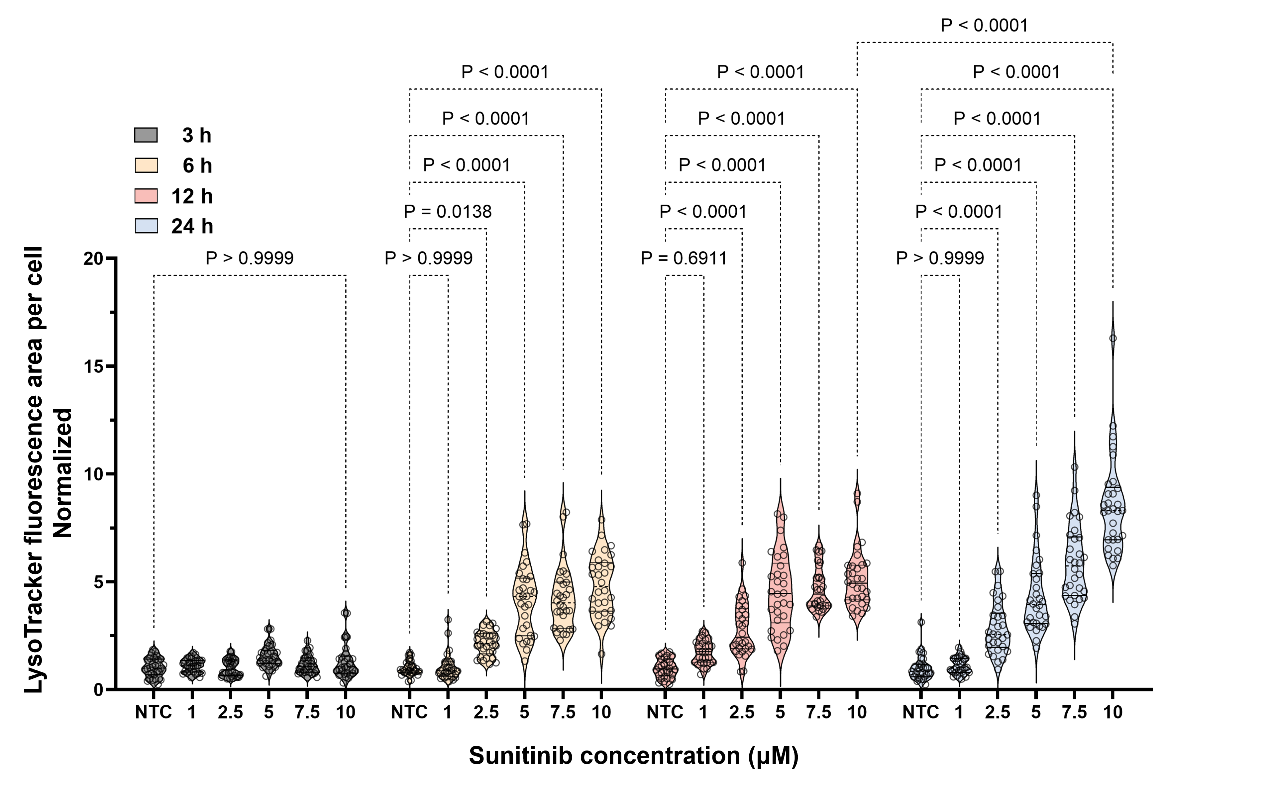

**Supplementary Fig. 5** Sunitinib concentration- and time-dependent LysoTracker™ Deep Red (LDR) fluorescence area per cell (normalized). All data was obtained by analyzing confocal microscopy images with ImageJ (fiji). Data are presented as violin plots, with the median and interquartile range indicated (N=3 independent experiments, n=10 cells analyzed per experiment, totaling 30 data points per group).

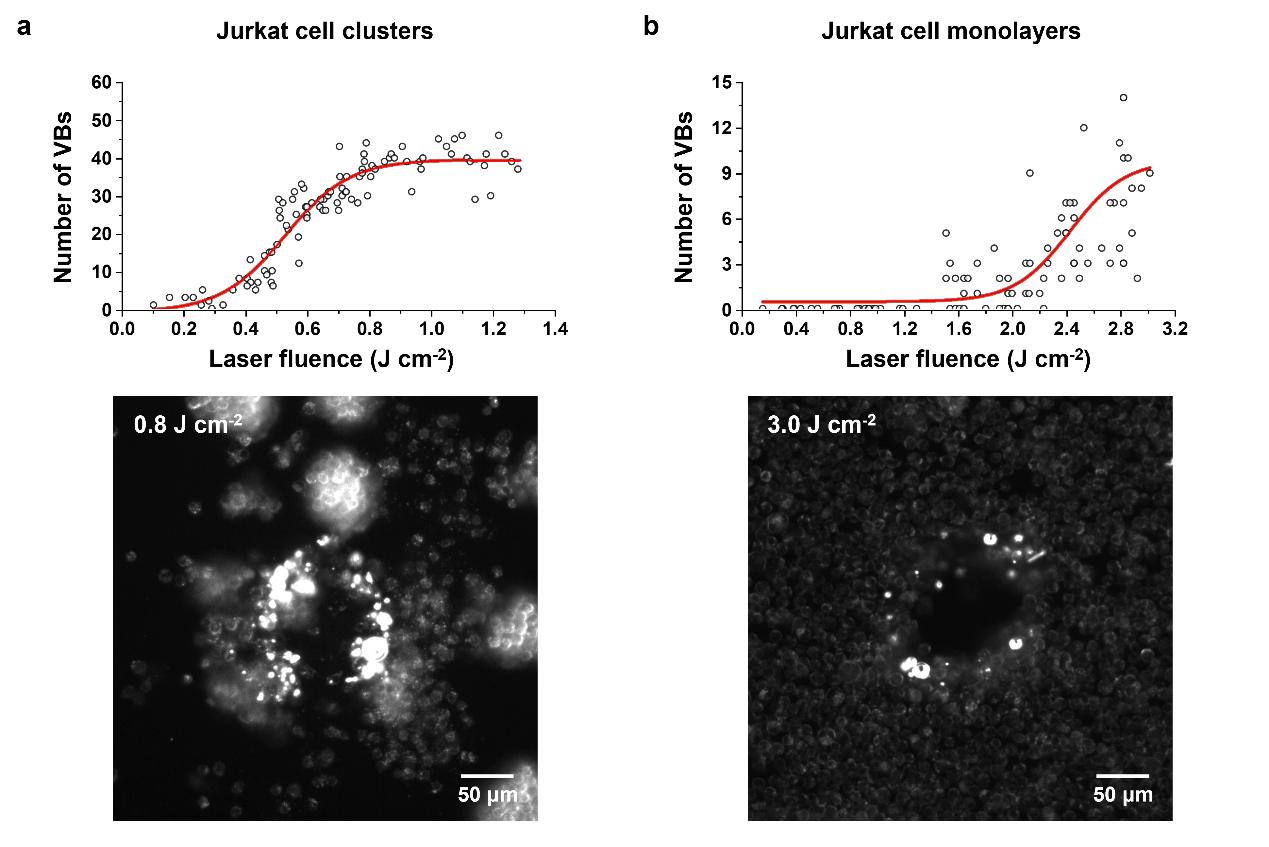

**Supplementary Fig. 6** Dark-field microscopy images of Jurkat cells incubated with 10 μM sunitinib, illustrating vapor bubble (VB) generation thresholds in two cellular configurations. **(a)** Jurkat cells spontaneously forming clusters. **(b)** Suspended Jurkat cells were briefly centrifuged (8 s pulse, 0-1000 rpm) to the bottom of a 96-well flat-bottom plate to create a uniform 2D cell monolayer. This configuration allows for a direct comparison of VB threshold dynamics between dense cellular clusters and cell monolayers under identical drug treatment conditions (10 μM sunitinib).

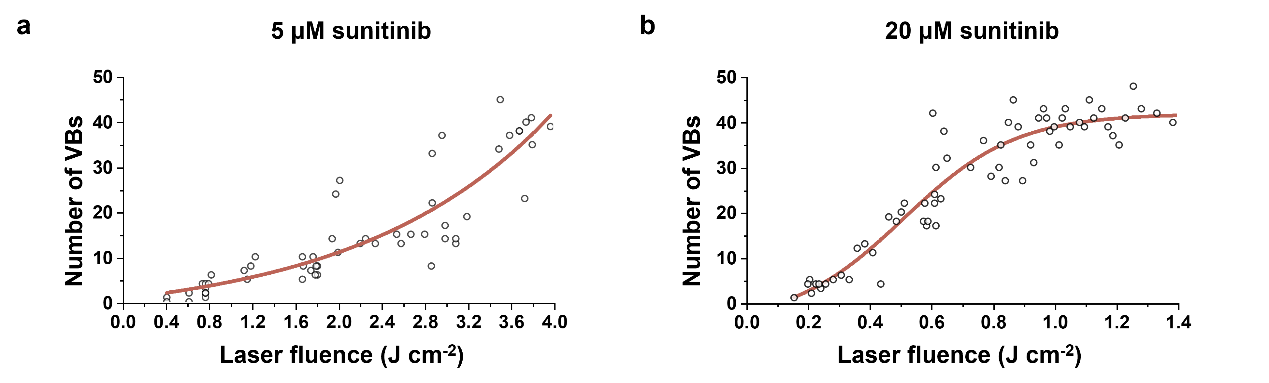

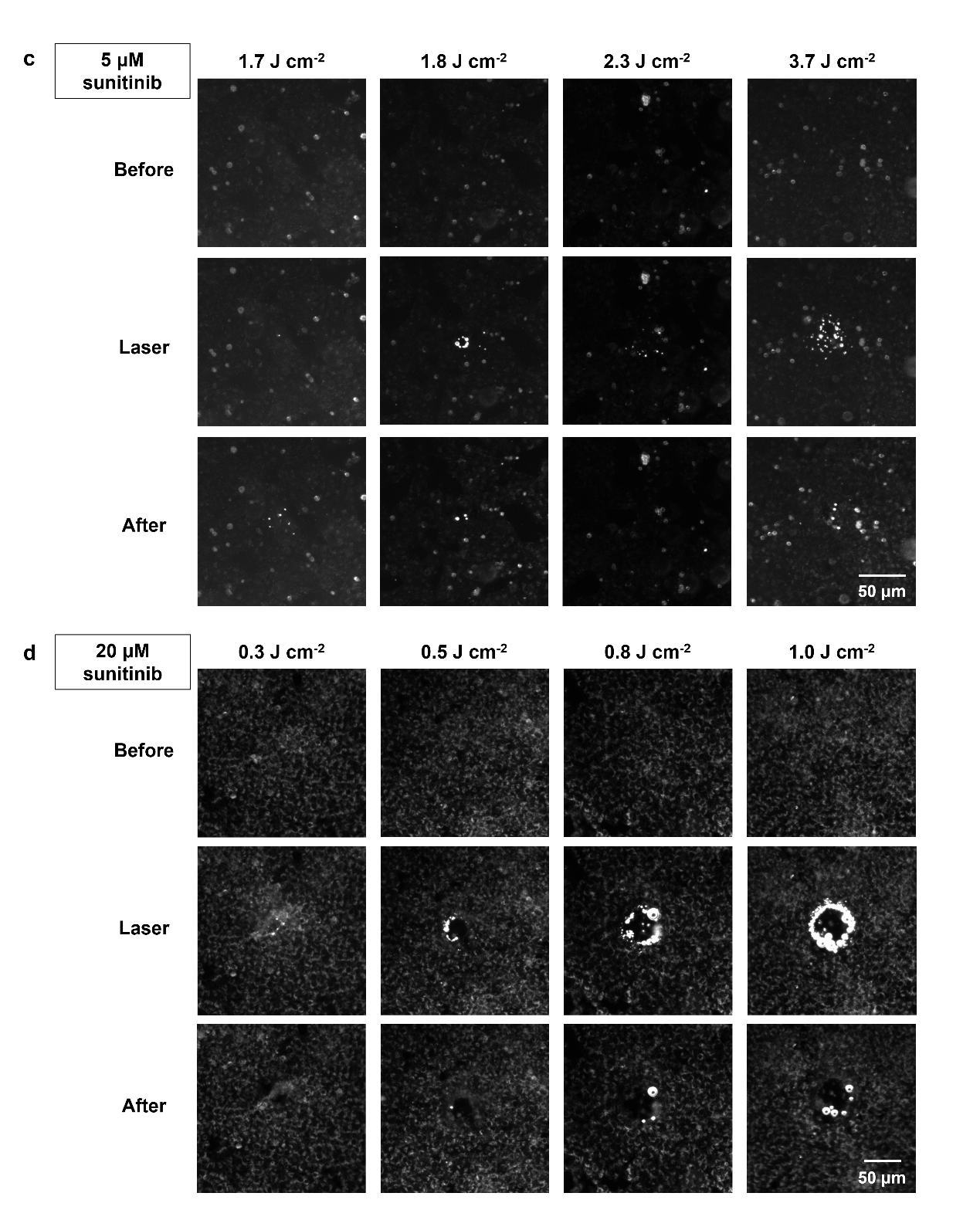

**Supplementary Fig. 7** Vapor bubble **(**VB) threshold for HeLa cells incubated with **(a)** 5 µM and **(b)** 20 µM sunitinib and corresponding dark field microscopy images of HeLa cells incubated with **(c)** 5 μM and **(d)** 20 μM of sunitinib before, during and after laser treatment with varying fluence.

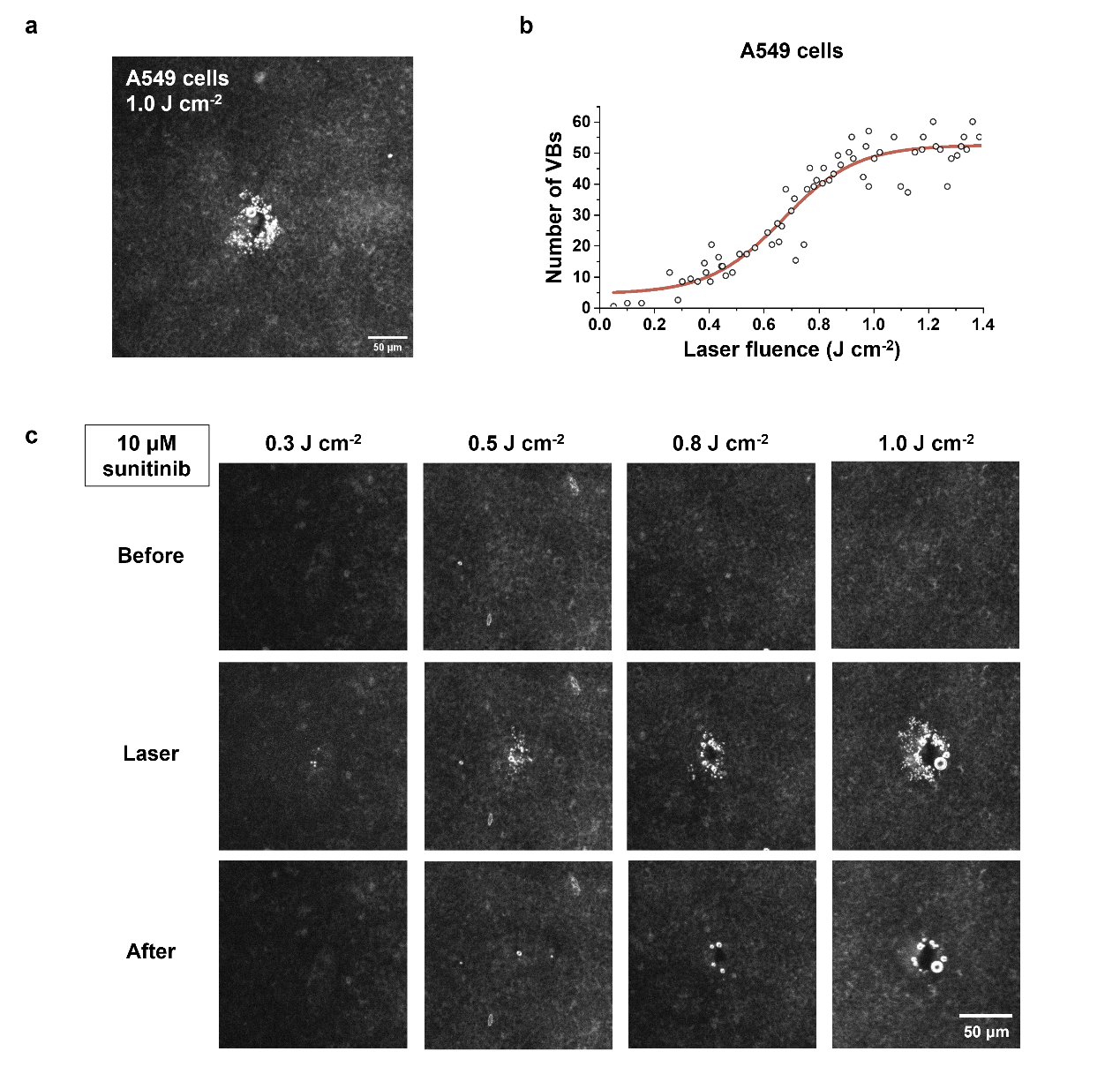

**Supplementary Fig. 8 (a)** Dark-field microscopy image of A549 cells incubated with 10 μM sunitinib and illuminated with a single 1.0 J cm^-2^ laser pulse. **(b)** Determination of the vapor bubble (VB) formation threshold. **(c)** Dark-field microscopy images of A549 cells incubated with 10 μM sunitinib before, during, and after laser irradiation at increasing laser fluence.

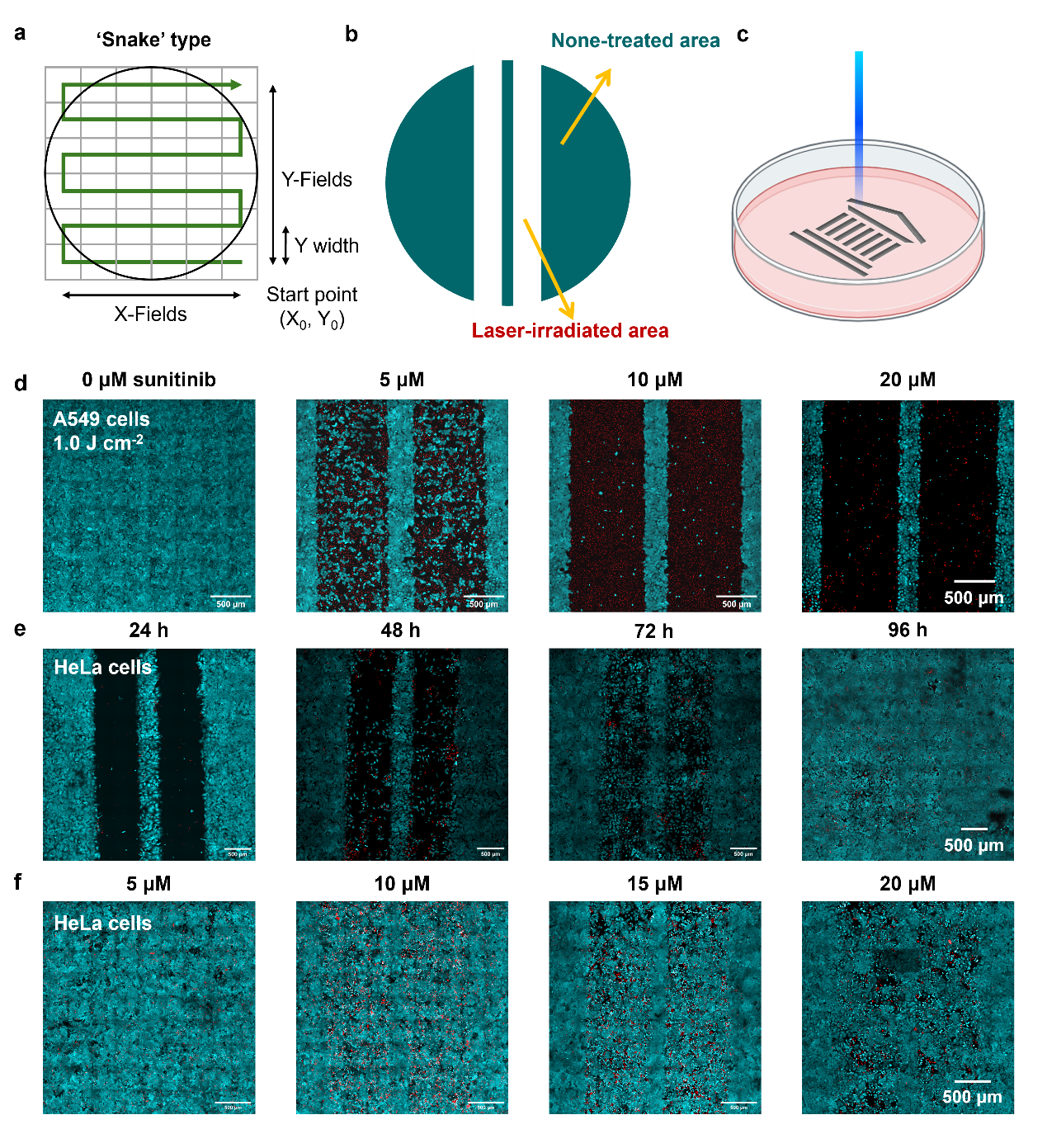

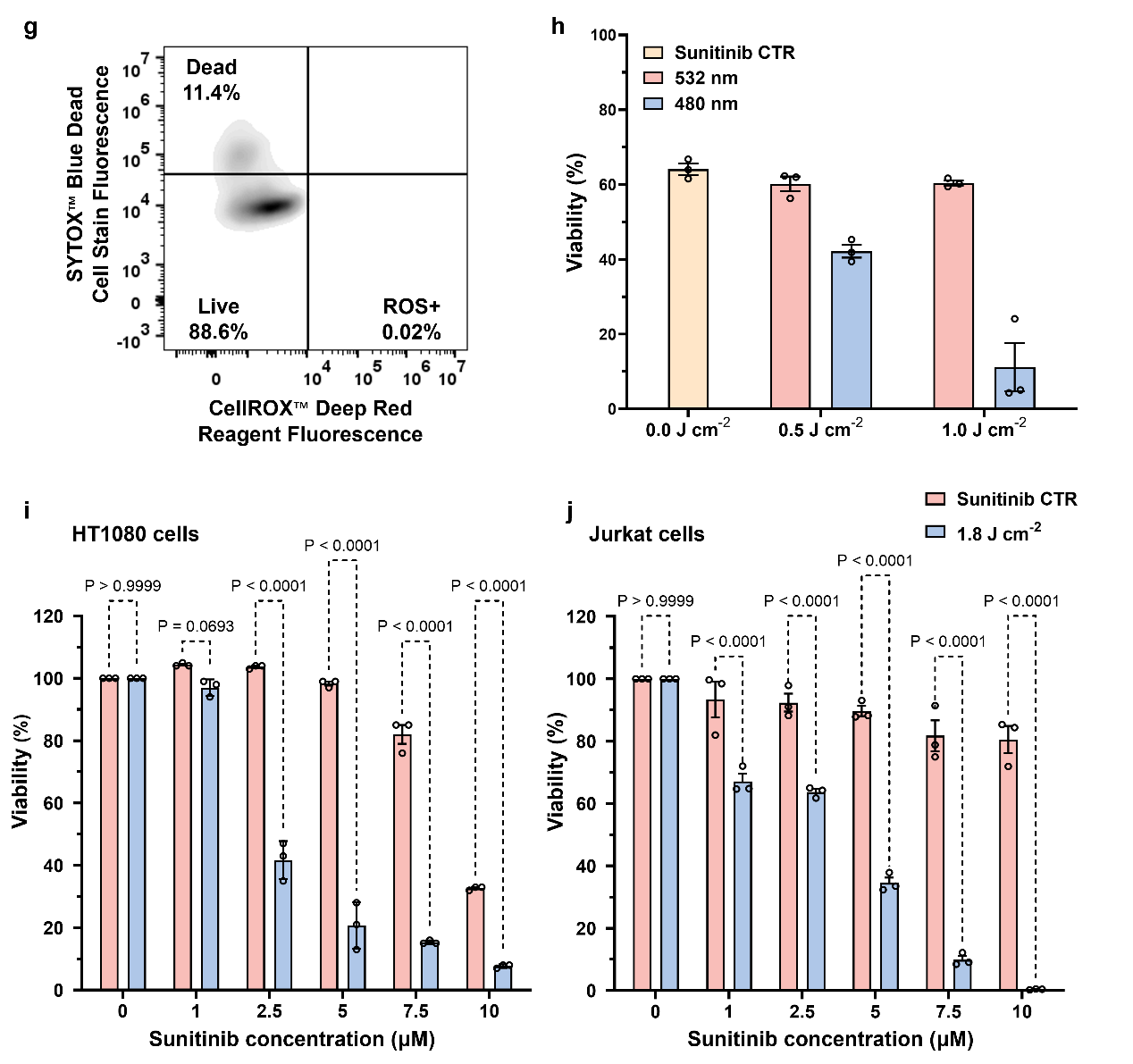

**Supplementary Fig. 9 (a-c)** Schematic of the applied pulsed laser scanning line and text pattern in cell killing experiments. **(d)** Confocal microcopy images of A549 cells treated with different concentrations of sunitinib (5-20 μM) using a fixed laser fluence (1.0 J cm^-2^). Green: Calcein-AM (live cells); Red: TO-PRO-3 (dead cells). **(e)** Images of HeLa cell recovery and outgrowth after laser irradiation. **(f)** Confocal microscopy images of laser treated Hela cells following sunitinib incubation (24 h) and sunitinib wash-out (72 h after medium replacement). **(g)** Flow cytometric analysis of intracellular ROS levels in Jurkat cells using CellROX™ Deep Red. The reagent was used in combination with SYTOX® Blue Dead Cell Stain to differentiate live stressed cells from dead cells. **(h)** Quantification of HeLa cell viability following laser treatment at different wavelengths (480 nm and 532 nm). Decrease in cell viability after laser treatment (1.8 J cm^-2^) applied on **(i)** HT1080 cells and **(j)** Jurkat cells incubated with sunitinib for 24 h at concentrations of 0-10 µM. Data are presented as mean ± SEM (N=3, n≥3).

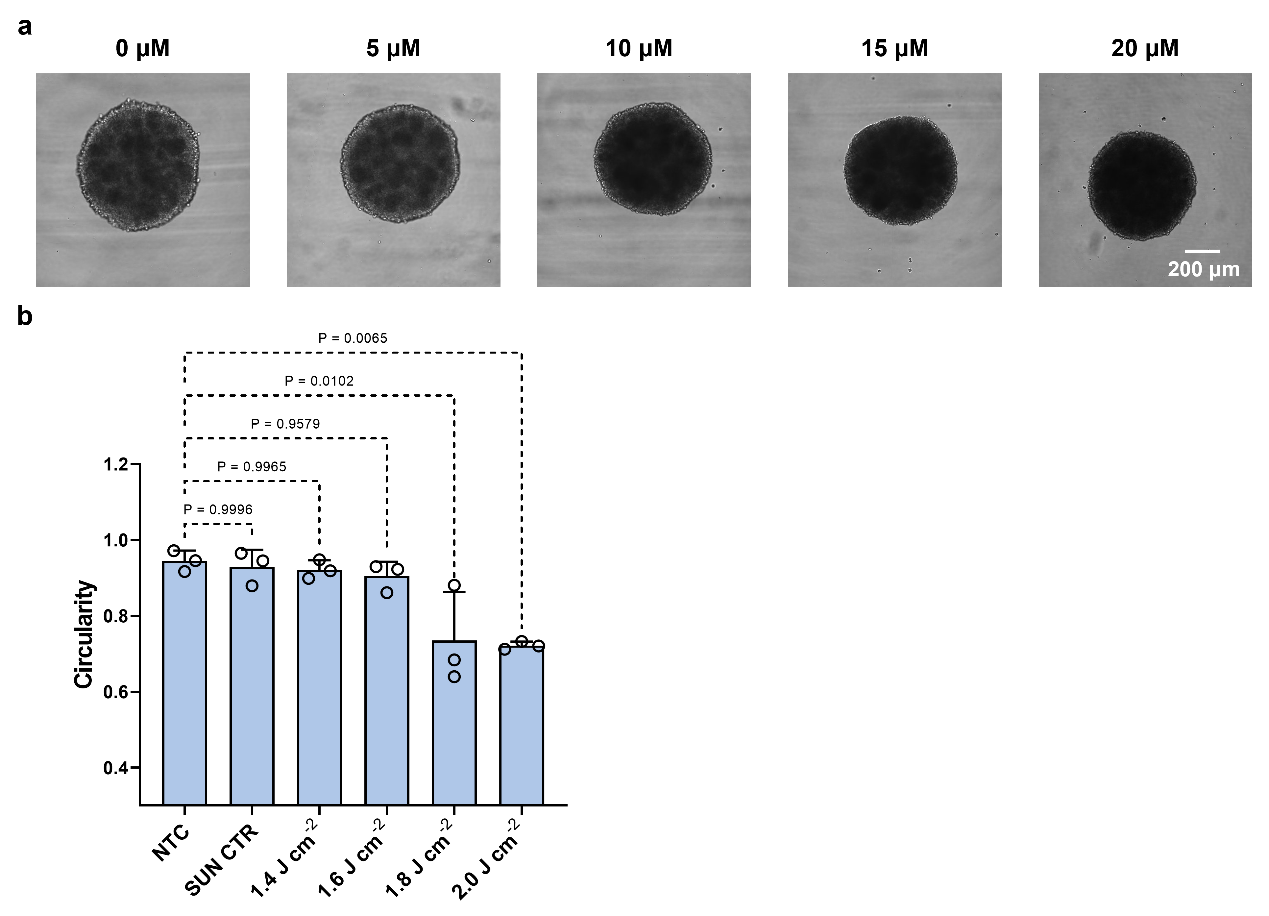

**Supplementary Fig. 10** (a) Optical microscopy images of HeLa spheroids incubated with increasing concentrations of sunitinib (0-20 µM) for 24 h. (b) Quantification of spheroid circularity. The reduction in circularity reflects structural disruption of spheroids induced by vapor bubble (VB) formation.

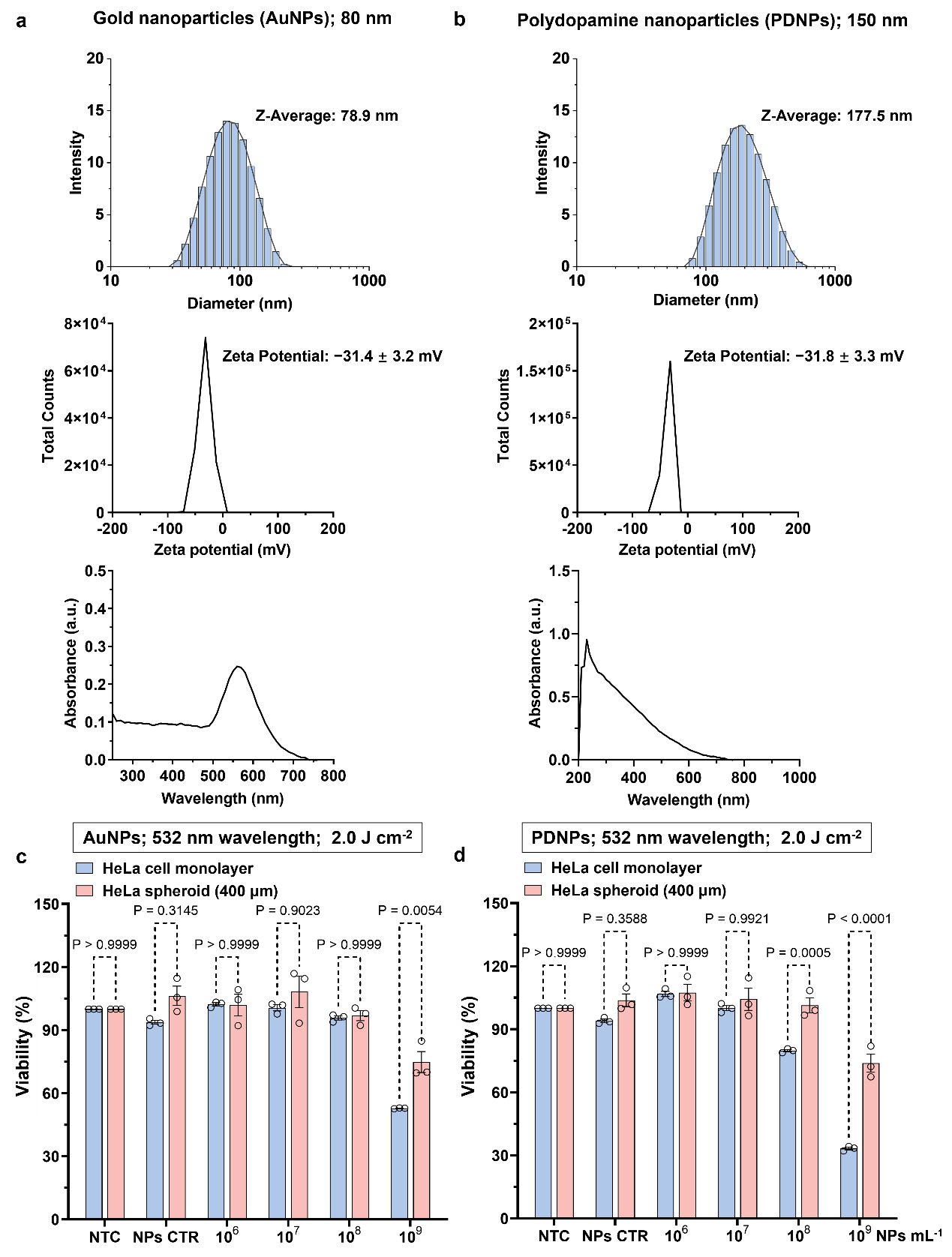

**Supplementary Fig. 11** Physicochemical characterization of **(a)** gold nanoparticles (AuNPs) and **(b)** poly(dopamine) nanoparticles (PDNPs). Size distribution (dynamic light scattering, DLS), UV-Vis absorption spectrum and zeta potential are shown. **(c)** Cell viability of HeLa cells in monolayer (2D) and spheroid (3D) cultures following 24 h incubation with 80 nm AuNPs and subsequent 532 nm pulsed laser irradiation. The marked difference in viability between 2D and 3D models highlights the limited penetration of AuNPs into the dense spheroid core. **(d)** Comparative cell viability analysis of HeLa cells in 2D and 3D cultures treated with 150 nm PDNPs for 24 h and 532 nm laser irradiation. The relatively high survival rate in the spheroid model reinforces the significant transport barriers and heterogeneous distribution of synthetic photothermal nanoparticles compared to small-molecular cationic amphiphilic drugs. Data are presented as mean ± SEM (N=3, n=3).

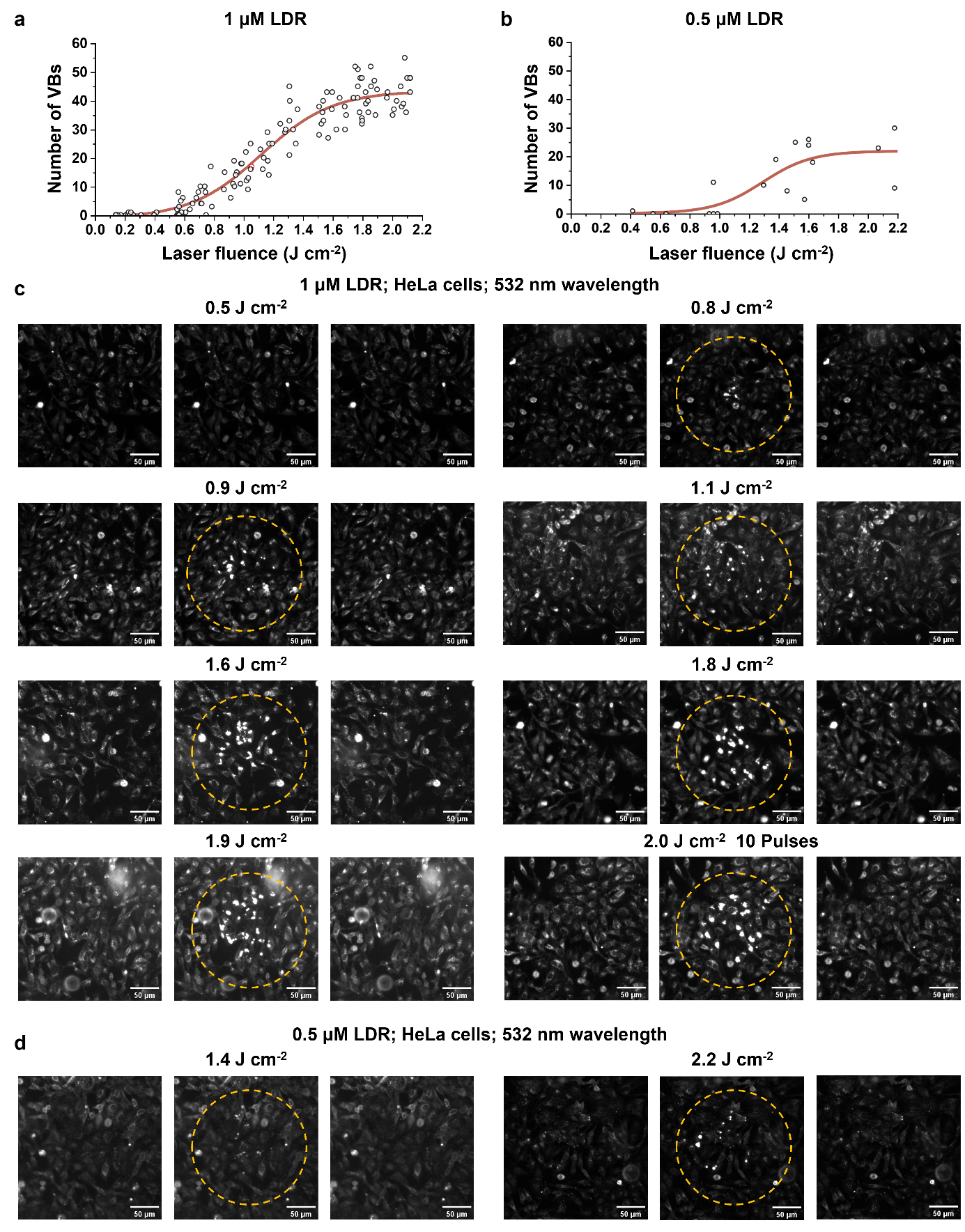

**Supplementary Fig. 12** Vapor bubble **(**VB) threshold for cells incubated with **(a)** 1 µM and **(b)** 0.5 µM Lysotracker™ Deep Red (LDR). **(c)** Corresponding dark field microscopy images of HeLa cells incubated with 1 µM LDR and **(d)** 0.5 μM LDR before, during and after 532 nm laser treatment with different laser fluence.

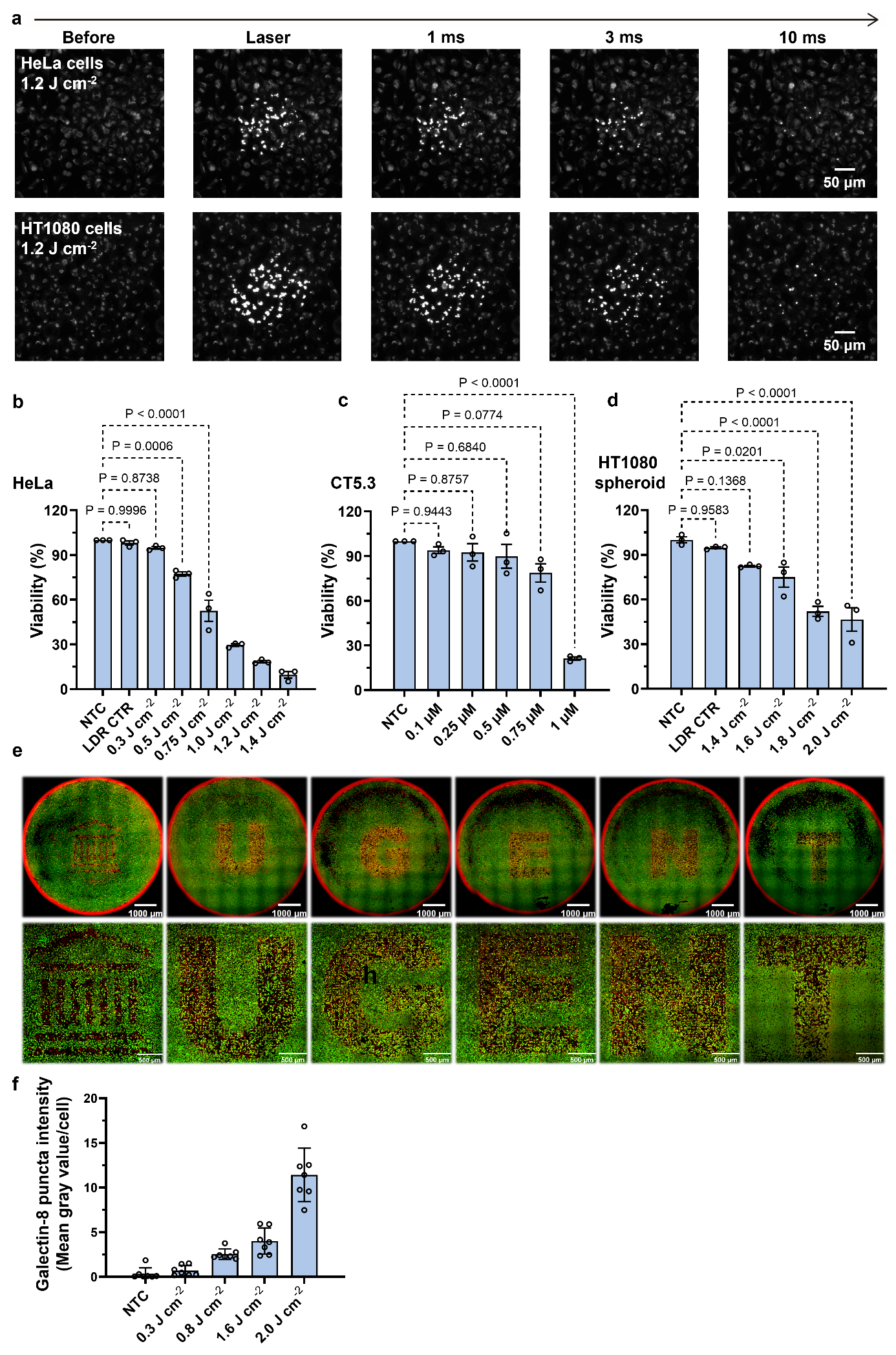

**Supplementary Fig. 13 (a)** Time-resolved dark-field microscopy images showing sequential frames of HeLa and HT1080 cells incubated with 1 μM Lysotracker™ Deep Red (LDR), before and after exposure to a 532 nm nanosecond pulsed laser (1.2 J cm^-2^). Vapor bubble (VB) generation and collapse events can be visualized in real time. **(b)** Quantification of HeLa cells viability following 24 h incubation with 1 μM LDR and subsequent irradiation at varying 532 nm laser fluences. **(c)** Viability of hTERT-immortalized CT5.3 cancer-associated fibroblasts (CAFs) incubated with varying concentration of LDR for 24 h and exposed to 1.2 J cm^-2^ laser pulses. **(d)** Quantification of HT1080 spheroids viability following laser treatment. Viability was assessed post-irradiation to evaluate the photomechanical impact of VB formation in LDR-treated spheroids. Data are presented as mean ± SEM (N=3, n=3). **(e)** Confocal fluorescence microscopy image showing spatially confined cell killing achieved via programmed laser scanning in a pre-defined text pattern. Green: Calcein-AM (live cells); Red: TO-PRO-3 (dead cells). **(f)** Quantification of GFP-Galectin-8 expression in MC38-eGFP-Galectin-8 cells following laser irradiation at different fluence levels. Data are presented as the mean gray value per cell, reflecting the degree of Galectin-8 recruitment under each irradiation condition. **(f)** Quantification of Galectin-8 recruitment into puncta within MC38-eGFP-Galectin-8 cells following laser irradiation at indicated fluences. Data are presented as the mean gray value of GFP-puncta per cell, reflecting the degree of lysosomal membrane permeabilization (LMP) under each condition. Data are presented as mean ± SEM (n=7). To quantify LMP-induced Galectin-8 recruitment, Z-stack images were first processed using the Maximum Intensity Projection (MIP) method within the Z-Project function of ImageJ (Fiji). This allowed the consolidation of high-intensity punctate structures from multiple focal planes into a single 2D projection, ensuring that all Galectin-8 recruitment sites throughout the cell volume were captured for analysis. To distinguish between the diffuse cytoplasmic GFP signal and the localized puncta, a fixed intensity threshold was applied to each projected image to isolate these high-intensity structures from the background. The diffuse green fluorescence in the cytoplasm was treated as a baseline and subtracted. The mean gray value of the detected puncta was then calculated per cell area, serving as a quantitative metric for the extent of Galectin-8 recruitment and, consequently, the degree of lysosomal damage triggered by vapor bubble (VB) formation.

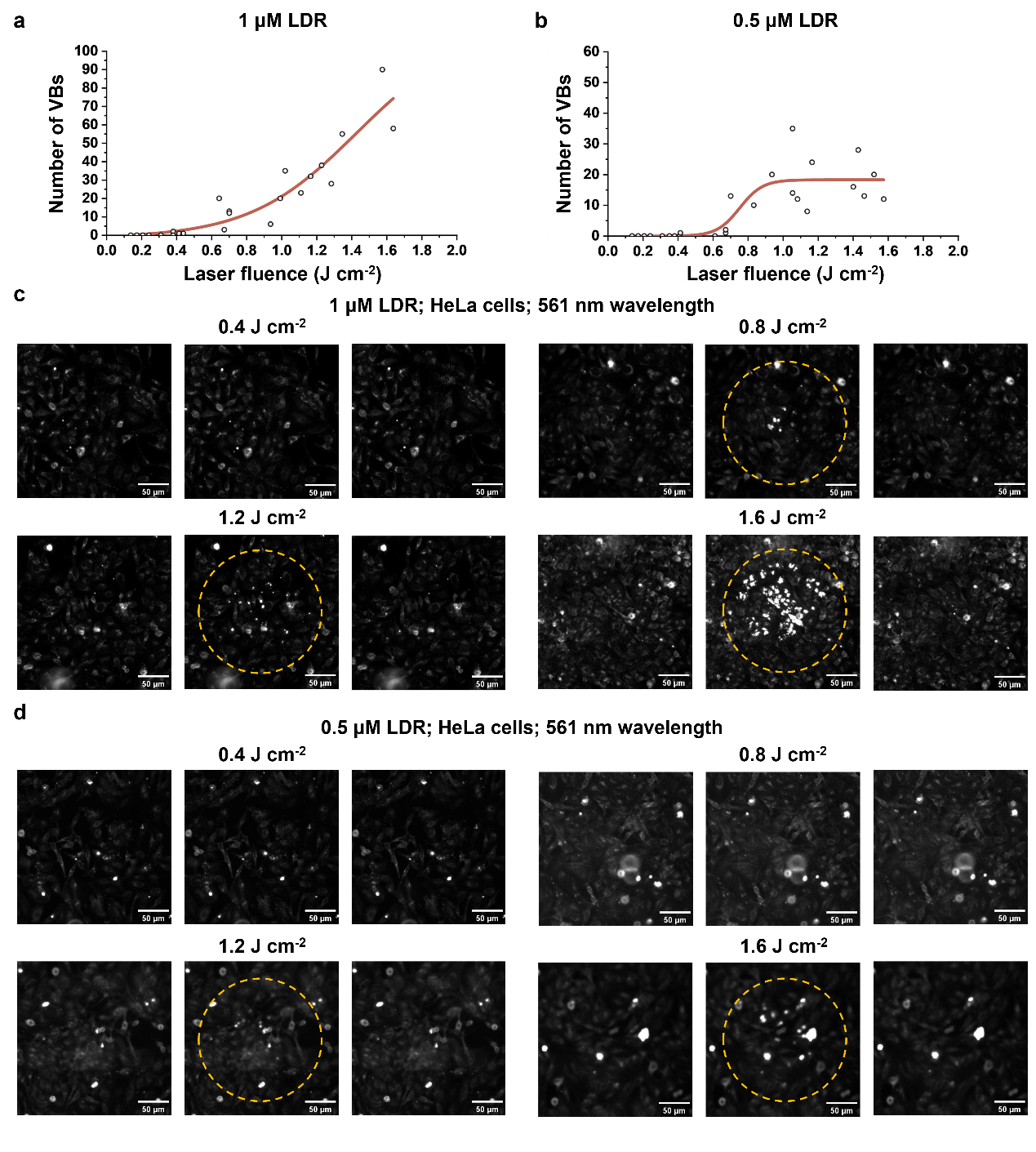

**Supplementary Fig. 14** Vapor bubble **(**VB**)** thresholds for cells incubated with **(a)** 1 µM and **(b)** 0.5 µM Lysotracker™ Deep Red (LDR) under 561 nm laser excitation. **(c)** Corresponding dark-field microscopy images of HeLa cells incubated with 1 µM LDR and **(d)** 0.5 µM LDR before, during, and after 561 nm laser irradiation at different laser fluence.

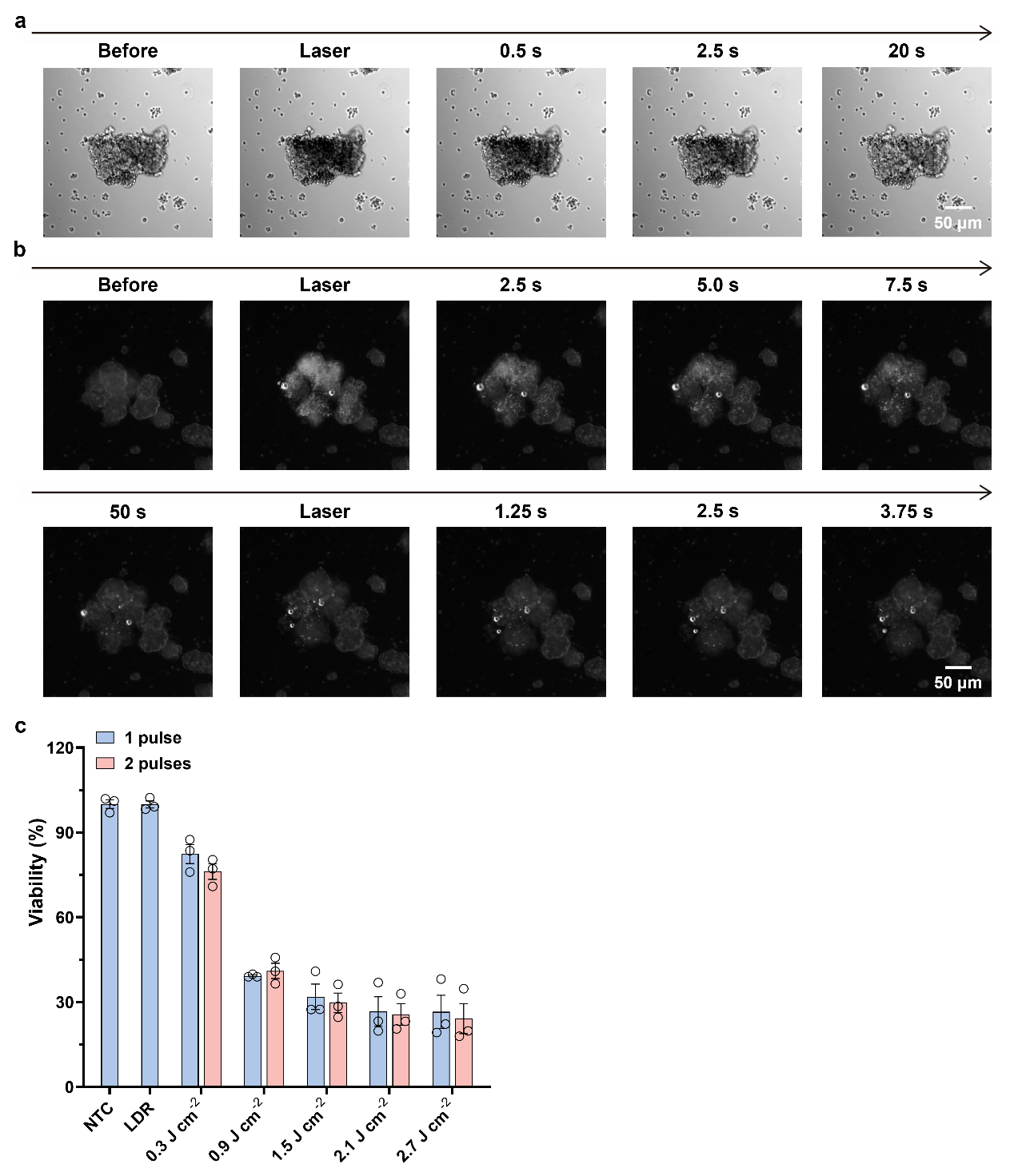

**Supplementary Fig. 15 Lysosomal vapor bubble-mediated tumor ablation in neuroblastoma tumoroids.** **(a)** Bright-field microscopy images of neuroblastoma tumoroids before and after nanosecond laser irradiation, showing a transient darkening at the tumor core immediately following a single pulse (2.0 J cm^-2^), indicative of vapor bubble (VB) formation and subsequent collapse. **(b)** Dark-field microscopy images of VBs in LysoTracker™ Deep Red (LDR)-incubated tumoroids upon laser exposure. A second laser pulse applied to the same region resulted in markedly reduced VB formation, likely due to lysosomal disruption after the first pulse. **(c)** Quantitative viability analysis of tumoroids treated with varying laser fluences (0.3-2.7 J cm^-2^), comparing single-pulse and two-pulse conditions. A significant viability drop was observed at and above the VB threshold (≥ 0.9 J cm^-2^), with no additional significant cytotoxicity upon repeated exposure, confirming that VB formation is the primary driver of photomechanical tumor ablation. Data are presented as mean ± SEM (N=3, n=6).
